# Dual transposon sequencing (Dual Tn-seq) to probe genome-wide genetic interactions

**DOI:** 10.1101/2024.09.24.614635

**Authors:** Justin J. Zik, Morgan N. Price, Adam P. Arkin, Adam M. Deutschbauer, Lok-To Sham

**Affiliations:** Infectious Diseases Translational Research Programme and Department of Microbiology and Immunology, Yong Loo Lin School of Medicine, National University of Singapore, Singapore; Environmental Genomics and Systems Biology Division, Lawrence Berkeley National Laboratory, Berkeley, CA 94720, USA; Department of Bioengineering, University of California, Berkeley, Berkeley, CA 94720, USA; Department of Plant and Microbial Biology, University of California, Berkeley, CA 94720, USA

**Keywords:** Functional genomics, *Streptococcus pneumoniae*, genetic interactions, CTP synthesis, peptidoglycan synthesis, nucleotide transport, transposon sequencing

## Abstract

Understanding gene function on a genome-wide scale is a fundamental goal in biology. With the advent of next-generation sequencing, high-throughput methods such as transposon sequencing (Tn- seq) were developed to measure fitness of single deletion mutants en masse. Nevertheless, gene redundancy complicates these approaches, as inactivating genes individually may not produce discernable phenotypes. Here, we report Dual Tn-seq, a technique for simultaneously assaying fitness of a comprehensive pool of double mutants. Dual Tn-seq couples random barcode transposon-site sequencing (RB Tn-seq) with the Cre-*lox* system, allowing us to determine the fitness of ≈68% of all possible double mutant combinations in the human pathogen *Streptococcus pneumoniae*. Genetic interactions identified from ≈1.4 billion double mutants uncovered new avenues in widely conserved biochemical pathways, exemplified by the discovery of a new CTP synthase and a regulator of peptidoglycan biosynthesis. Since Dual Tn-seq has very few requirements, it can be readily adapted to diverse organisms.

## INTRODUCTION

Knowing the sequence of a gene does not necessarily imply understanding its function. Even in the best-studied bacterium, *Escherichia coli*, about one-third of its genes encode “proteins of unknown function” and “hypothetical genes.” Traditional approaches for elucidating gene function, such as phenotyping deletion mutants, are relatively time-consuming and labor-intensive. Furthermore, single- knockout mutants may not have any detectable phenotype due to gene redundancy. Because our ability to sequence genomes outpaces functional characterization, most genes are annotated solely based on sequence and structural similarities ^1^. While powerful tools like AlphaFold emerged recently ^2^, they have yet to achieve accuracies to replace experimental validations ^1^. Additionally, having a predicted structure alone does not necessarily clarify function. The limited understanding of gene function precludes the development of pathway-directed chemical screens ^3^, which may contribute to a largely unexplored space of drug targets. Moreover, a subset of hypothetical genes in pathogenic bacteria like *Streptococcus pneumoniae* differs from those in *E. coli*, further illustrating the need for an effective functional genomic approach that can be applied across multiple species.

One approach to uncovering gene function is constructing a loss-of-function mutant and profiling its transcriptome, proteome, metabolome, and phenome ^4,5^. Yet, this multiomics approach is unsuitable for studying numerous hypothetical genes, especially because some are functionally redundant ^6^. Another approach is to leverage genetic interactions to infer gene function. A genetic interaction occurs when the phenotype of a double mutant deviates from expectation based on the phenotypes of the two single mutants. A classic example is synthetic lethality, where the individual mutants in two genes have no or minimal fitness defects, but the double mutant has a severe fitness defect. As genes that form such relationships are often functionally related, we can use this information to guide mechanistic studies. It also partially overcomes the problem of gene redundancy. For example, genetic interaction arrays such as GIANT-coli ^7^, E-MAP ^8^, and SGA ^9^ were developed. These methods directly quantify the fitness of many double mutants by systematically measuring colony sizes or growth in liquid cultures ^10,11^. Nevertheless, the prerequisites of these approaches include an extensive collection of single knockout mutants and robotics, which may not be available for most researchers. Consequently, a complete genetic interaction array is only available in yeast ^12^.

By contrast, transposon insertion sequencing (TIS) approaches (e.g., Tn-seq ^13^, TraDIS ^14^, and IN-seq ^15,16^) assay individual gene knockouts without requiring preconstructed single mutants for every gene. Here, a transposon (Tn) mutant library is generated from a strain of choice. The location of Tn insertions and their abundance are determined by next-generation sequencing of the Tn junctions, and hence fitness of most genes in the genome can be assayed in parallel. To discover genetic interactions, the results of a Tn mutant library constructed in the wild-type background versus a mutant background can be compared ^17–19^ . A rate-limiting step of TIS is the preparation of the DNA library for sequencing the genomic DNA adjacent to the Tn insertions. To increase the throughput of TIS, random barcode transposon-site sequencing (RB Tn-seq) was developed by inserting a unique DNA barcode within the Tn ^20^. After associating the insertion sites and barcodes by TIS, the linked barcodes serve as a surrogate for mapping and quantifying the Tn mutants via barcode sequencing (BarSeq).

While RB Tn-seq and traditional TIS are powerful for genetic interaction studies, they are “one-to-all” approaches in which any gene(s) under study must first be mutated prior to Tn library construction. By design, traditional TIS approaches cannot identify two random Tn insertions in a single cell, as these are almost exclusively beyond sequencing distance for individual templates. To overcome these challenges, we developed dual transposon sequencing (Dual Tn-seq). Dual Tn-seq is designed for an “all-to-all” genome-wide mining of genetic interactions by measuring the fitness of a massive number of double mutants in parallel. We chose the human pathogen *Streptococcus pneumoniae* as our prototype because it is naturally competent and is genetically tractable ^21^. Briefly, two barcoded Tn libraries were generated in *S. pneumoniae* strain D39W. Genomic DNA was purified from one of the libraries to transform the other, generating ∼1.4 billion double mutants. To identify and quantify the double mutants, the Tns have an engineered *lox* site so that the barcodes would be recombined when Cre was induced, and thus bring them within the distance for Illumina sequencing, regardless of the orientation of Tn insertions. After sequencing ∼10 billion of the fused barcodes, the fitness of each double mutant was determined by comparing the expected and the actual number of reads. As a result, we could reliably sample approximately 68% of the total 1.3 million possible double mutant combinations. The screen identified ∼6,000 genetic interactions, some of which were confirmed by independently reconstructing the double mutants. As expected, these pairs are often mapped to the same biochemical pathway, illustrating the power of Dual Tn-seq in uncovering gene functions. For instance, we discovered a new regulator of peptidoglycan synthesis and unraveled a novel CTP synthase. Additionally, the nucleoside transporters in *S. pneumoniae* were identified and validated. As Dual Tn-seq relies on only a few components, it can be adapted to elucidate the “genetic dark matter” in a wide range of bacterial species.

## RESULTS

### Design, construction, and validation of barcoded Tn libraries for dual Tn-seq

We constructed two HIMAR1 mariner Tns with spectinomycin or erythromycin resistance cassettes (RBloxSpec and RBloxErm). They contain sequencing primer binding sites, a *lox66* or *lox71* site, and a 20-nucleotide barcode (**Fig. S1A**). These Tns are expected to recombine to produce a dual barcode sequence when *cre* expression is induced. The barcode fusion is irreversible because the *lox66* and *lox71* sites will generate a non-functional *lox72* site after recombination ^22^.

Tn insertions are expected to occur in random orientations. Yet, the *lox* sites are directional during recombination. We reason that if the *lox* sites in the Tns are placed as shown in **Fig. S1**, the barcodes will be fused regardless of the orientation of Tn insertions. Upon *cre* induction, the chromosome between the insertions will be excised or inverted, depending on the orientation of the two Tns (**Fig. S1 and S2A)**, thereby fusing the barcodes. To demonstrate this, we constructed two strains in which RBloxSpec was inserted at penicillin-binding protein 1b (*pbp1b* or *spd*_*1925*), followed by introducing RBloxErm at penicillin-binding protein 1a (*pbp1a* or *spd*_*0336*) in either orientation. *pbp1a* and *pbp1b* are 493 kb apart. Nevertheless, we readily detected barcode fusion in cells harboring both RBlox-Tns in either orientation when *cre* was induced (**Fig. S2B and S2C**). One concern is that the basal expression level of *cre* may lead to growth defects in the double mutant because it may excise or invert the chromosome. Yet, we did not detect differences in the growth rates of the Δ*pbp1a* Δ*pbp1b* mutants in the presence or absence of the *cre* expression cassette (P_Zn_-*cre*) (**Fig. S2D**). This result suggests the slower growth in the Δ*pbp1a* Δ*pbp1b* mutant was solely due to the inactivation of penicillin-binding proteins. We conclude that the Cre-*lox* system can recombine the genome specifically to fuse the barcodes inserted in different chromosomal locations and orientations.

Next, we constructed two plasmid libraries from the RBloxSpec and RBloxErm cassettes. Vectors harboring each cassette were barcoded and introduced into *E. coli*, generating pools with 15.2 million and 8.8 million unique barcodes for RBloxSpec and RBloxErm, respectively. Plasmids purified from each pooled library were mixed with MarC9 transposase and genomic DNA of *S. pneumoniae* for *in vitro* transposition. After end-repairing and purification, the transposed DNA was introduced into the prototype pneumococcal strain D39W harboring an inducible *cre*. The Tn insertions and the associated barcodes of each pool were sequenced and mapped (**Fig. S3A**), revealing hundreds of thousands of insertions with unique barcodes per library (see **methods**). Collectively, these libraries covered insertions within the central part (10-90%) of the open reading frame in ∼1,600 of the 1,914 protein-coding genes of the pneumococcal genome.

As a proof-of-concept, we recapitulated known synthetic lethal relationships between *pbp1a* and *pbp2a* (*spd_1821*) and *pbp1a* and *macP* (*spd_0876*) ^23–25^ using the barcoded libraries. To do this, we transformed the wild-type and Δ*pbp1a* strains with the RBloxErm library and sequenced the barcodes in the transformants (**Fig. S3B**). As expected, there were fewer Tn insertions in *pbp2a* or *macP* when the library was constructed in the Δ*pbp1a* background compared to the wild-type (**Fig. S3C**). Additionally, insertions in *pbp1a* were markedly reduced when we transformed the RBloxErm library into Δ*pbp2a* and Δ*macP* mutants (**Fig. S3C**), indicating that the Bar-seq approach can reproduce results from traditional Tn-seq screens.

### Dual Tn-seq identifies genetic interactions in *S. pneumoniae*

Under standard laboratory growth conditions, *S. pneumoniae* has ∼1,600 non-essential genes ^13,26,27^. Theoretically, at least 1.3 million combinations of double mutants can be made. To approach saturation of the screen, we first optimized the transformation protocol (**Fig. S4A and S4B**). Next, we transformed the RBloxErm library with DNA purified from the RBloxSpec library, resulting in ∼1.4 billion double mutants (**Fig. 1A**). The double mutants were pooled and allowed to recover for two hours (∼1 doubling) before *cre* induction. Cells were further incubated for two hours before harvesting. Quantitative PCR (qPCR) showed that at this time point, the barcode recombination had reached saturation (**Fig. S4C**). Genomic DNA was extracted, and dual barcodes were amplified and sequenced, generating over 10 billion reads.

**Fig. 1.**
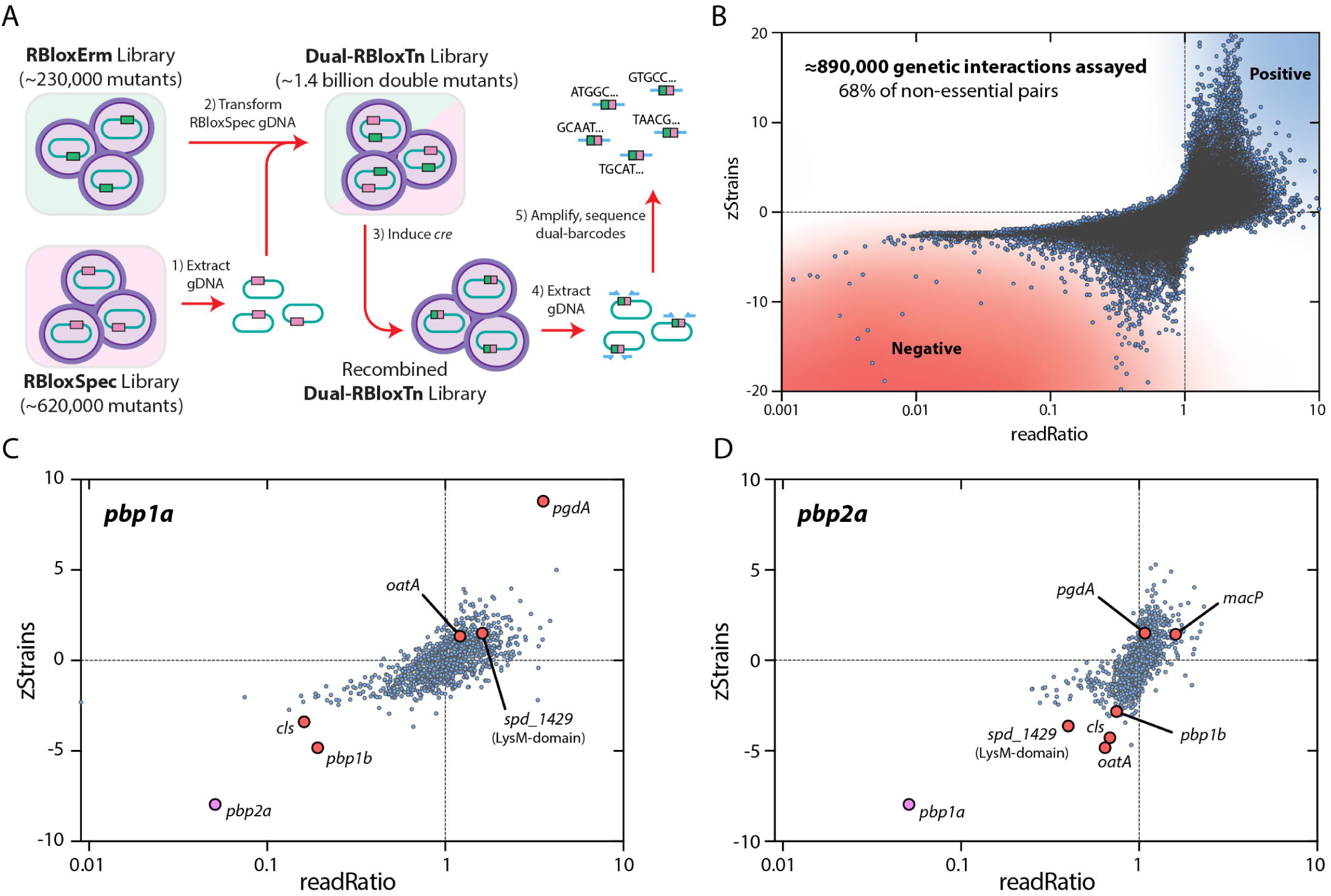
Dual Tn-Seq identifies genetic interactions in *S. pneumoniae*. (**A**) The overall workflow. Two DNA-barcoded Tn mutant libraries (RBloxErm and RBloxSpec) were constructed, each containing hundreds of thousands of unique mutants. Genome DNA was extracted from the RBloxSpec library and used to transform the RBloxErm library, generating ∼1.4 billion double Tn mutants. Next, *cre* was induced to recombine the DNA barcodes. DNA was extracted from the Dual RBlox Tn library, and the fused barcodes were amplified and sequenced. (**B**) In total, 889,918 gene pairs were examined by Dual Tn-seq, covering 68% of all double mutants that can be theoretically constructed. readRatios are plotted at the detection threshold for clarity (i.e. observed reads + 1 for all). A lower ZStrains value and readRatio correlate with negative genetic interactions (**Fig. S5**). 13 interactions are outside the current graph limits. Representative Dual Tn-seq results, exemplified by *pbp1a* (**C**) and *pbp2a* (**D**). As expected, *pbp1a* and *pbp2a* (in pink) showed a strong negative genetic interaction. Other genes of interest, mainly in the peptidoglycan synthesis pathway, are highlighted in red. Data points on the y-axis have a readRatio value of 0.

We used two metrics to identify pairs of genes with significant genetic interactions, both negative interactions (covering both synthetic lethality where the double mutant is completely lethal and synthetic sick where the double mutant is less fit but still viable) and positive interactions (**Fig. S5**). The first metric, ‘readRatio,’ is calculated by dividing the number of observed dual-barcode reads for a given gene pair by the expected number, determined via a probabilistic model from the abundances of individual gene barcodes (**Fig. S5**). The second metric, ’zStrains,’ uses the expected number of double mutant strains (from unique barcode combinations) as the mean of a Poisson distribution to calculate a Z-score describing the distance, in standard deviations, of the observed number of strains from expectation (**Fig. S5**). These metrics are affected globally but uniformly by the location and distance between the two Tn insertions across the chromosome (**Fig. S6**), allowing us to correct this bias mathematically (**Table S1**). Moreover, gene pairs with too few reads or too close to each other were filtered (see **methods**). To better visualize the putative genetic interactions, we plotted the readRatios against the zStrains (**Fig. 1B**). As expected, *pbp1a* and *pbp2a* are among the negative genetic interactions identified by the dual Tn-seq (**Fig. 1C and 1D**). We also confirmed a negative genetic interaction between *pbp1a* and *pbp1b* described previously^23^ (**Fig. 1C and S2D**). However, *pbp1a* and *macP* could not detected as a synthetic lethal pair because of insufficient coverage, likely because of the small size of the *macP* gene. Together, 889,918 non-essential gene pairs (68%) have sufficient coverage, which could be used for mining genetic interactions.

Next, we sought to empirically determine values of read ratio and zStrains that represent bona fide negative genetic interactions, both synthetic lethal and synthetic sick. We selected 109 random gene combinations that spanned a wide range of these metrics and asked whether or not we could construct these double mutants by transforming a knockout construct of one gene into the corresponding deletion background. A drastic decrease in transformation efficiency or small colony sizes would indicate negative genetic interactions. In addition, we transformed an unrelated deletion cassette to control for the transformation efficiency (**Fig. S7A and Table S2**). With a stringent threshold of a 100-fold reduction of transformation efficiency in our assay, we found 15 synthetic lethal gene pairs in this pool (**Fig. S7B**). Five additional double mutants formed small colonies on the plate, suggesting they are synthetically sick (**Fig. S7B and S7C**). As expected, we observed a strong correlation between synthetic lethality and decreased zStrains and readRatios (**Fig. S7D and S7E**). Thus, we ranked the gene pairs using both metrics to prioritize candidates for mechanistic studies. Next, we used a shuffle test and functional relatedness to choose thresholds with a low rate of false positives (see Methods). At a false discovery rate of ∼7%, we identified 86 negative genetic interactions (**Table S3**). Our thresholds include most (15/20) of the gene pairs that we validated as synthetic lethal or as yielding small colonies. Nevertheless, some confirmed synthetic lethal gene pairs (e.g., *murZ* and *yjbK*) are outside this threshold, suggesting that the number of genetic interactions identified by Dual Tn-seq is larger.

### *spd_0310* encodes a novel ammonium-dependent CTP synthase

We identified a strong negative interaction between *pyrG* (*spd_0442*) and *spd_0310* (**Fig. 2A and 2B**). PyrG is the only known CTP synthase in all life forms ^28,29^. It produces CTP by amination of UTP, primarily using L-glutamine as the amine donor (**Fig. S8**) ^30^. Spd_0310 contains a DUF1846 domain and has been linked to iron homeostasis and virulence, though its molecular function remains unknown ^31^. To confirm this genetic interaction, we placed *spd_0310* downstream of the zinc-inducible promoter. A significant growth defect was observed when *spd_0310* was depleted in a Δ*pyrG* background (NUS4797; Δ*spd_0310* Δ*pyrG* // P_Zn_-*spd_0310*) (**Fig. 2C and 2D**). To test if *spd_0310* participates in CTP biosynthesis or salvage, we examined whether exogenous cytidine could rescue the growth defect in NUS4797. Indeed, cytidine supplementation corrected the defect, while adding other nucleosides did not (**Fig. 2C, S9A, and S9B**).

**Fig. 2.**
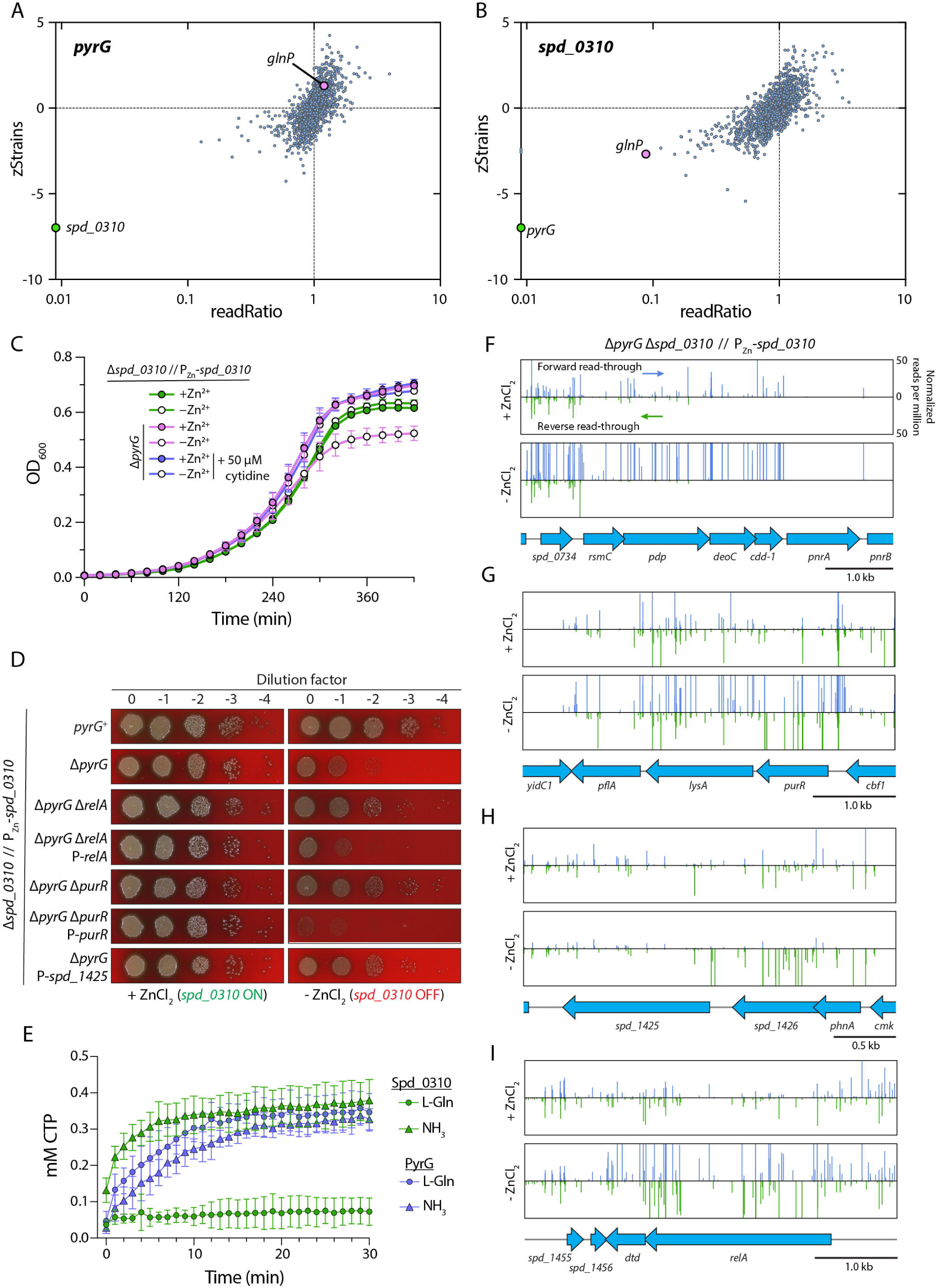
PyrJ (Spd_0310) is a CTP synthase. (**A**) *pyrG* exhibited a strong negative genetic interaction with *pyrJ* (in green). (**B**) *pyrJ* likely formed a synthetic lethal pair with *pyrG* (in green) and showed a negative genetic interaction with *glnP* (in pink), which encodes a glutamate permease. Data points on the y-axis have a readRatio value of 0. (**C**) Strains NUS3924 [Δ*spd_0310* // P_Zn_-*spd_0310*] and NUS4797 [Δ*pyrG*(*spd_0442*) Δ*spd_0310* // P_Zn_-*spd_0310*] were grown in BHI medium at 37°C in 5% CO_2_ with added ZnCl_2_ and MnCl_2_, washed twice, and resuspended in BHI medium with or without cytidine and/or ZnCl_2_/MnCl_2_ supplementation. Growth was monitored by measuring the optical densities of the cultures at 600nm (OD_600_) over time. Plotted are the means of three biological replicates and the standard deviation. (**D**) The indicated strains (**Table S4**) were grown in BHI broth at 37°C in 5% CO_2_ until the OD_600_ was between 0.2 and 0.4. Cultures were normalized based on their optical density, serially diluted, and spotted on blood agar plates with (left) or without (right) added ZnCl_2_ and MnCl_2_. Plates were incubated overnight at 37°C in 5% CO_2_ before imaging. Genes following “P-” are overexpressed from a constitutive promoter. Experiments were done twice with similar results. (**E**) Purified PyrJ (Spd_0310) converted UTP to CTP using ammonia but not glutamine as the amine donor. By comparison, PyrG used glutamine or ammonia for CTP synthesis. Plotted are the means and standard deviation from three independent experiments. (**F to I**) RB Tn-seq revealed the nucleoside transporters in pneumococcus. An RB Tn library was constructed by transforming strain NUS4797 with the genomic DNA from a mapped library (RBloxErm). At least one million transformants were obtained. Transformants were collected and plated on blood agar with or without added ZnCl_2_ and MnCl_2_. Plates were incubated overnight at 37°C in 5% CO_2_ before the cells were harvested for barcode sequencing. The number of reads per million of transposon insertions at the positive (forward read-through) or negative (reverse read-through) strand was enumerated. The genomic loci of *pnrAB* (**F**), *purR* (**G**), *spd*_*1425* (**H**), and *relA* (**I**) are shown.

Next, we performed RB Tn-seq in NUS4797 to identify mutations that suppress the lethality of Δ*spd_0310* Δ*pyrG*. When *spd_0310* was depleted in a Δ*pyrG* background, there was an increase in insertions upstream of *pnrABCD* and *spd_1425*. PnrABCD is a nucleoside transporter, while Spd_1425 is an uncharacterized major facilitator superfamily transporter. Notably, the insertions are enriched only in the same orientation as the transcription of *pnrABCD* and *spd_1425* (**Fig. 2F, 2H**). As the Tn has no terminator, it is likely the Tn insertions led to the overexpression of these transporters. Expectedly, overexpression of *spd_1425* alleviates the growth defects (**Fig. 2D**), suggesting that both transporters are likely involved in nucleoside import, thereby relieving the nucleotide starvation caused by the deficiencies of PyrG and Spd_0310. To confirm this speculation, we showed that inactivating *spd_1425* and *pnrABCD* exacerbates the phenotype of Δ*pyrG* upon *spd_0310* depletion (**Fig. S9C**). In addition, Δ*spd_1425* and Δ*pnrABCD* abolished cytidine uptake, as judged by the lack of rescue by adding exogenous cytidine to NUS4797 (**Fig. S9D**).

The screen also revealed the upstream regulatory pathways of CTP synthesis. Insertions disrupting the *pur*-operon repressor *purR* (*spd_1776*) and the stringent response gene *relA* (*spd_1458*) also suppressed the growth defect of NUS4797 when *spd_0310* was not depleted (**Fig. 2G and 2I**). PurR is implicated in controlling purine/pyrimidine biosynthesis ^30,32^, and thus Δ*purR* may increase the pool of CTP in the cell. On the other hand, RelA induces the stringent response by synthesizing the alarmone (p)ppGpp. Since (p)ppGpp is an anti-inducer of PurR ^33^, a *relA* mutation will increase the level of nucleotide synthesis by de-repressing PurR.

To test whether Spd_0310 is involved in nucleoside uptake, we exposed the mutants to the toxic analog 5-fluorocytidine (5FC) ^34^. Cells unable to salvage cytidine are resistant to 5FC, while those with increased uptake should be hypersensitive. Confirming the RB Tn-seq results, both Δ*purR* and Δ*relA* mutants showed reduced growth in the presence of 5FC compared to the parent strain (**Fig. S9E**). Ectopic expression of *spd_1425* inhibited growth under 5FC, indicating enhanced cytidine uptake. Conversely, Δ*spd_1425*, but not Δ*pnrABCD*, drastically reduced 5FC sensitivity, suggesting that Spd_1425 is the primary cytidine transporter in *S. pneumoniae*. These results are consistent with the finding that PnrA prefers purine nucleosides ^35^.

Next, we constructed a Δ*spd*_*1425* Δ*pnrABCD* double mutant to test if cytidine uptake was abolished. Indeed, the double mutant was resistant to 5FC, but it exhibited a slightly reduced growth rate compared to the Δ*spd_1425* single mutant (**Fig. S9E**). Combining Δ*spd*_*1425* Δ*pnrABCD* with Δ*pyrG* or Δ*spd_0310* did not affect 5FC sensitivity, as both triple mutants behaved similarly to the parent strains (**Fig. S9F**). These findings strongly suggest that Spd_0310 is involved in CTP synthesis rather than nucleoside transport, and confirm that Spd_1425 is a cytidine transporter. Finally, we purified Spd_0310 and demonstrated that it converts UTP to CTP. Unlike PyrG, which can use glutamine or ammonia as amine donors, Spd_0310 exclusively uses ammonia (**Fig. 2E and S10**). Consistently, Δ*spd_0310* has a negative genetic interaction with the glutamine transporter gene *glnP* (**Fig. 2B**). Cells lacking *glnP* likely have a reduced glutamine pool in the cell and decrease PyrG activity (**Fig. 3A**). Given its role as an alternative CTP synthase, we renamed *spd_0310* as *pyrJ*.

**Fig. 3.**
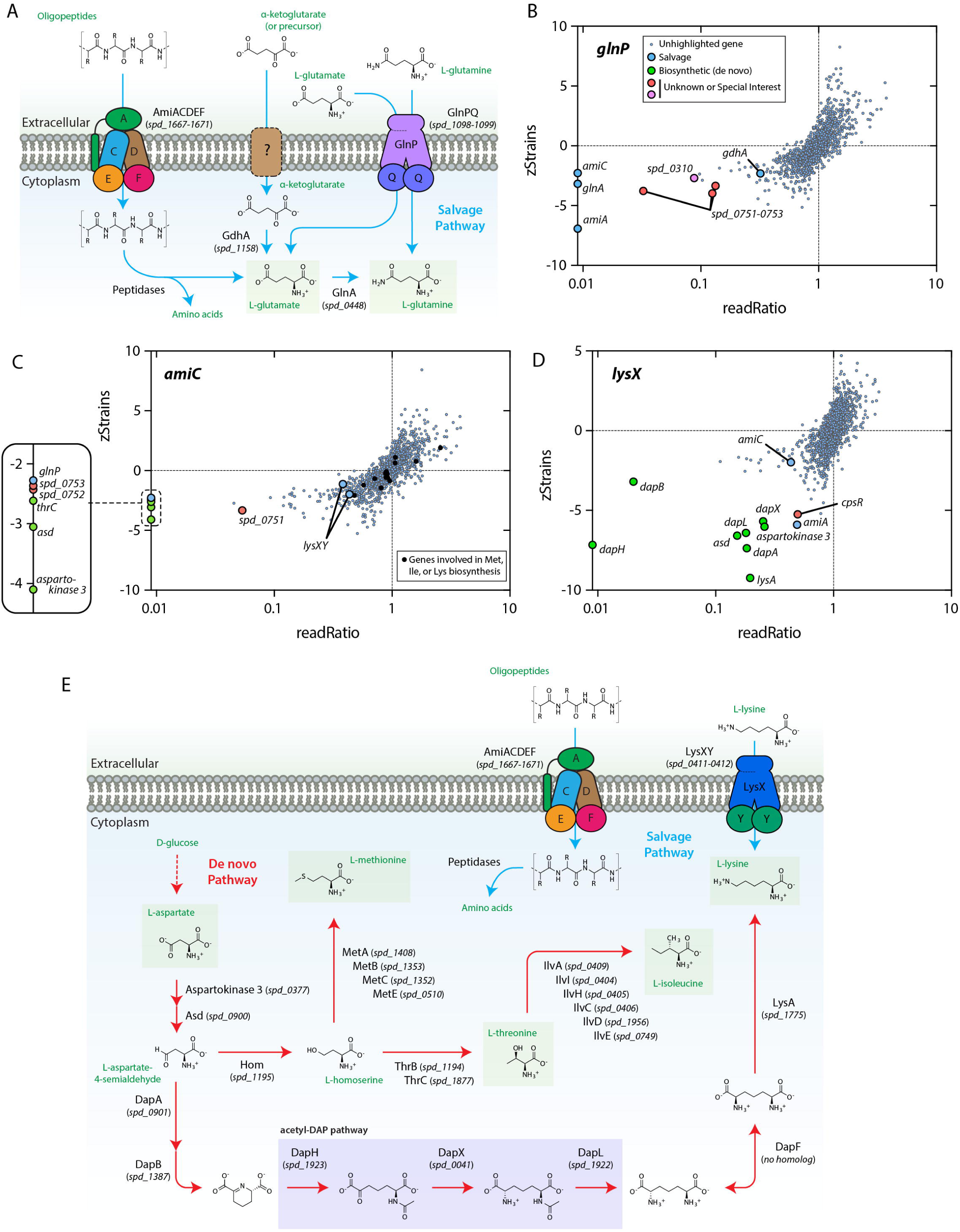
Genetic interactions identified amongst amino acid salvage and biosynthesis pathways. (**A**) Glutamine is imported via the GlnPQ transporter (Spd_1098-1099) or synthesized by GlnA (Spd_0448) using glutamate as a precursor. Additionally, GdhA (Spd_1158) can generate glutamate from _J-ketoglutarate, which is likely imported into the cell. The Ami transporter complex (Spd_1667- 1671; AmiACDEF) salvages peptides from the extracellular environment, providing an additional source of glutamate. (**B**) *glnP* shows negative interactions with the glutamine synthase gene *glnA and* other genes that presumably contribute to the glutamine pool. Data points on the y-axis have a readRatio value of 0. (**C**) The oligopeptide transporter gene *amiC* exhibits strong negative interactions with several amino acid biosynthesis genes and other known or presumed amino acid transporter systems. The black dots represent the genes that are exclusively involved in methionine, isoleucine, and lysine synthesis. Data points on the y-axis have a readRatio value of 0. (**D**) The lysine transporter gene *lysX* interacts negatively with genes in the *de novo* lysine biosynthesis pathway. Data points on the y-axis have a readRatio value of 0. (**E**) The lysine biosynthetic and salvage pathways in *S. pneumoniae*. Lysine biosynthesis branches from methionine, threonine, and isoleucine biosynthesis at L-asparatate-4-semialdehyde.

### Redundant transporters revealed by Dual Tn-seq

Transporters are often required for virulence as they acquire essential nutrients from the host ^36^. *S. pneumoniae* encodes two Trk/Ktr/HKT family potassium transporters (CabP/TrkH and Spd_0429/Spd_0430). Whether strains lacking both systems could survive remains controversial ^18,37^, possibly because of the difference in the growth media. Trk/Ktr/HKT family transporters often comprise a transmembrane component and a positive regulator ^38^. For example, CabP (Spd_0077) contains a regulator of conductance of K^+^ (RCK) domain and activates TrkH (Spd_0076). When bound to cyclic-di-AMP (c-di-AMP), CabP interaction with TrkH is reduced, causing a decrease in potassium uptake ^37^. We found that *trkH* and c*abP* form synthetic lethal pairs with *spd*_*0429* and *spd*_*0430*, suggesting that one of these potassium systems is required for survival *in vitro* (**Fig. S11A and S11B**). Spd_0429 is the transmembrane component and requires the regulatory protein Spd_0430 for function. Notably, Spd_0430 has two RCK domains, and thus, it may sense additional ligands such as NAD^+^, NADH, or Ca^2+^ ^38^. In summary, either potassium acquisition system can support growth under our experimental conditions, and both regulatory proteins are required for the function of their cognate transmembrane channel.

Similarly, *S. pneumoniae* encodes two ATP-binding cassette (ABC) transporters for inorganic orthophosphate (P_i_): Pst1 and Pst2 ^39^. These are encoded by the *pst1* operon (*spd_1910*-*spd_1914*) or the *pst2* operon (*spd_1227*-*spd_1232*). Downstream of these operons are genes encoding the negative regulatory proteins PhoU1 and PhoU2. PstA and PstC form the transmembrane channel, which interacts with the cytoplasmic ATPase PstB and the solute-binding protein PstS. We detected strong negative genetic interactions between genes of the two Pst systems (**Fig. S11C and S11D**), consistent with their redundant roles in P_i_ transport. Unexpectedly, despite their inhibitory roles in their cognate systems, fewer insertions in *phoU1* and *phoU2* were also detected in *pst2* and *pst1* Tn mutants, respectively. This is likely due to polar effects of the Tn insertions, as strains lacking both *phoU* homologs have reduced viability ^39^. Thus, Dual Tn-seq is capable of revealing components of multi-protein complexes and pathways.

### Genetic interactions identified between the salvage and *de novo* biosynthesis pathways

Like the redundant transporters, we detected strong genetic interactions between genes encoding *de novo* biosynthetic enzymes and factors in the corresponding salvage pathway. For example, *ribU* (*spd_0437*) encodes the small integral membrane subunit (S-component) of the energy-coupling factor (ECF)-type ABC transporter ^40^. S-components are modular, membrane-embedded substrate-binding proteins that interact with a shared ECF-transporter to dictate its specificity. Expectedly, the riboflavin-specific *ribU* has negative genetic interactions with riboflavin synthesis genes (*ribBA*, *ribDG*, *ribH*, or *ribE*) (**Fig. 4**).

**Fig. 4.**
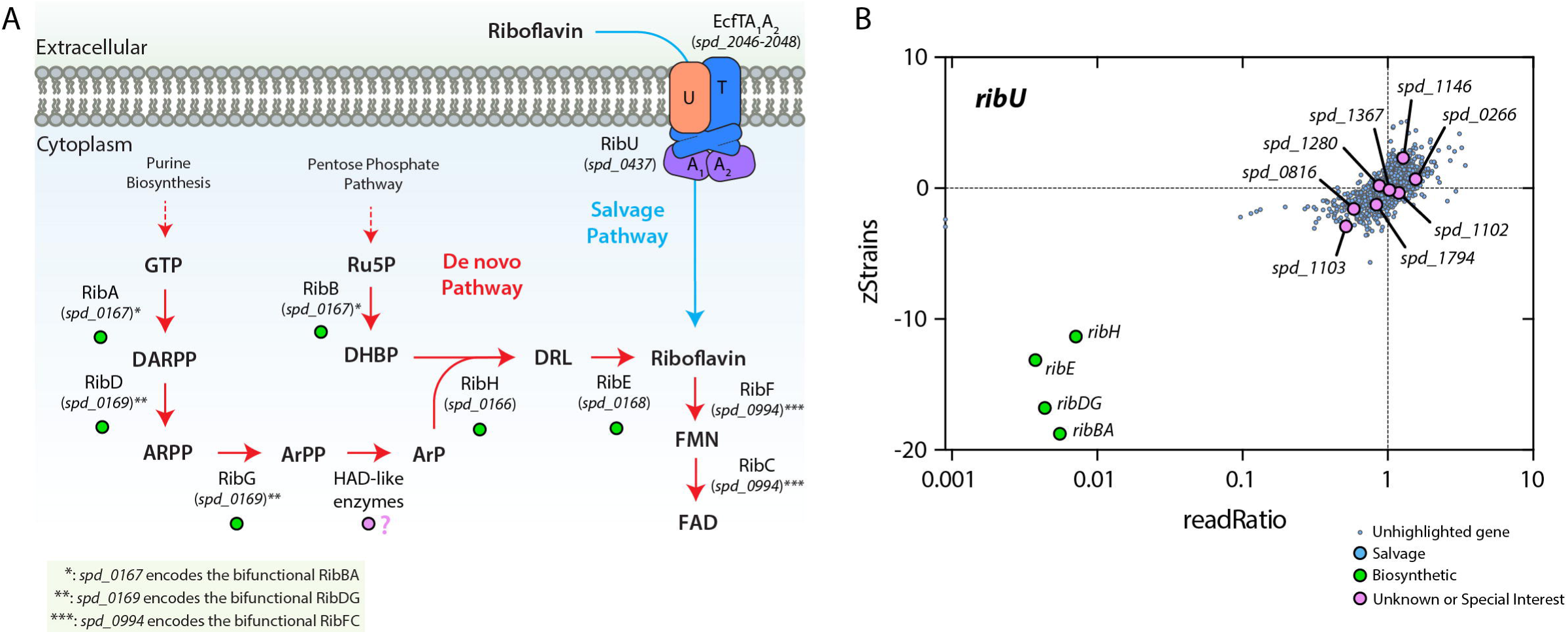
Genetic interactions identified between riboflavin salvage and *de novo* biosynthesis pathways. (**A**) The riboflavin biosynthetic and salvage pathways in *S. pneumoniae*. GTP, guanosine 5’-triphosphate; DARPP, 2,5-diamino-6-ribosylamino-4(3H)-pyrimidinone 5’-phosphate; ARPP, 5- amino-6-(ribosylamino)-2,4-(1H,3H)-pyrimidinedione 5’-phosphate; ArPP, 5-amino-6-ribitylamino- 2,4(1H,3H)-pyrimidinedione 5’-phosphate; ArP, 5-amino-6-ribitylamino-2,4(1H,3H)-pyrimidinedione; Ru5P, D-ribulose 5-phosphate; DHBP, 3,4-dihydroxy-2-butanone-4-phosphate; DRL, 6,7-dimethyl-8- (1-D-ribityl)lumazine; FMN, flavin mononucleotide; FAD, flavin adenine dinucleotide. (**B**) The riboflavin transporter gene *ribU* negatively interacts with genes in the riboflavin biosynthetic pathway (green data points). Purple data points show the eight homologs of the ArPP phosphatases in *B. subtilis*. We did not detect strong negative genetic interactions between these genes and *ribU*, likely because they are functionally redundant.

Riboflavin is synthesized from the products of purine biosynthesis and the pentose phosphate pathway (**Fig. 4A**). At the intersection of these pathways, RibH converts 5-amino-6-ribitylamino- 2,4(1H,3H)pyrimidinedione (ArP) to 6,7-dimethyl-8-(1-D-ribityl)lumazine (DRL), which is converted to riboflavin by RibE (**Fig. 4A**). ArP is produced by dephosphorylating 5-amino-6-ribitylamino- 2,4(1H,3H)pyrimidinedione 5’-phosphate (ArPP). We did not detect a strong candidate for ArPP phosphatase by Dual Tn-seq. Recently, proteins in the haloacid dehalogenase (HAD) superfamily have been implicated in dephosphorylating ArPP ^41^. As *S. pneumoniae* has eight homologs of the ArPP phosphatases identified in *Bacillus subtilis*, it is likely some are redundant here as in other organisms^42^ (**Fig. 4**).

De novo synthesis of the active form of vitamin B6 (pyridoxal phosphate (PLP)) is catalyzed by the PdxST complex (*spd_1297*-*spd_1296*) (**Fig. S12A**). PLP can also be obtained via a salvage pathway (**Fig. S12A**). Here, pyridoxal (PL), pyridoxine (PN), or pyridoxamine (PM) are imported and subsequently converted to PLP. PL can be directly phosphorylated by PdxK to generate PLP, while PN and PM need to be oxidized after phosphorylation ^43^. The PNP/PMP oxidase that mediates this step remains unknown in pneumococcus ^43^. We detect strong negative genetic interactions between PdxST and the known salvage pathway components (PdxU2, PdxK) (**Fig. S12B, S12C, and S12D**). PdxU2 is an ECF-transporter S-component specific for PL, PN, and PM ^44^. The screen also reported negative interactions between the salvage pathway and PdxR (Spd*_*1225) (**Fig. S12B, S12C, and S12E**), a transcription activator that stimulates *pdxST* expression ^45^, illustrating that Dual Tn-seq can uncover factors in the whole biochemical pathway.

### Dual Tn-seq uncovers candidate factors for amino acid synthesis

Many bacteria synthesize glutamate and glutamine from the Krebs (TCA) cycle intermediate _J- ketoglutarate. Since *S. pneumoniae* lacks the Krebs cycle, it is a glutamate and glutamine auxotroph^46,47^ (**Fig. 3A**). Yet, it maintains the enzyme (GdhA, Spd_1158) that converts _J- ketoglutarate to glutamate, suggesting it can salvage exogenous _J-ketoglutarate (**Fig. 3A**). The ABC transporter GlnPQ (*spd_1098-spd_1099*) mediates the import of glutamine and glutamate. As expected, it has negative genetic interactions with the glutamine synthase gene *glnA* (*spd_0448*) and *gdhA* (**Fig. 3**). Additionally, the oligopeptide ABC transporter Ami (*spd_1667-spd_1671*) likely supports glutamate/glutamine acquisition, as suggested by the negative interactions with *glnP*. A three-gene operon (*spd_0751-spd_0753*) was also revealed as having negative interactions with *glnP* and *ami* (**Fig. 3B and 3C**). Consistently, simultaneous deletion of *amiE* and *spd_0751* resulted in a small colony phenotype (**Fig. S7C**). Spd_0751 has remote structural homology to PTS transporters and arsenate pumps. Spd_0752 resembles a gluconate permease from a Foldseek search. Finally, Spd_0753 is likely a pyrrolidone-carboxylate peptidase (PCP), a widespread peptidase that cleaves L- pyroglutamyl residues (a derivative of glutamate) at the N-terminus of peptides ^48^. Thus, we speculate that *spd_0751-spd_0753* is an alternative peptide transport system or an _J-ketoglutarate importer. Consistently, *spd_0751-spd_0753* expression increases in a Δ*glnA* Δ*glnP* double mutant ^49^.

We observed strong negative interactions between the lysine transporter genes *lysXY* (*spd_0411- spd_0412*) and genes in the *de novo* lysine biosynthesis pathway (**Fig. 3D and 3E**) ^50^. Although *S. pneumoniae* synthesizes lysine using the acetyl-diaminopimelic acid (acetyl-DAP) pathway, it lacks *dapF* for the diaminopimelate epimerase step ^46,50^. Despite the selective pressure and the importance of lysine synthesis in the Δ*lysX* mutant, our analysis did not identify a diaminopimelate epimerase, with the only other similarly clustered gene being *cpsR* (*spd_0064*), a transcriptional regulator presumably controlling capsule production ^51^. This suggests that the diaminopimelate epimerase enzyme might be redundant, essential for growth, or below our detection threshold. Nevertheless, these findings support the annotation of genes in this pathway and demonstrate the efficacy of Dual Tn-seq in gene function elucidation.

As expected, the interactions between transporter genes (*ami* and *lysXY*) were less pronounced because the cells can rely on *de novo* lysine biosynthesis (**Fig. 3D**). However, strong negative interactions were detected between *ami* genes and genes for the early steps of lysine synthesis, aspartokinase 3 (*spd_0377*) and *asd* (*spd_0900*). Asd produces L-aspartate-4-semialdehyde, which is at the branch point between lysine and methionine/threonine/isoleucine biosynthesis (**Fig. 3E**). Inactivation of *asd* and aspartokinase 3 also abolishes *de novo* threonine synthesis. We detected a negative genetic interaction between *ami* and *thrC*, suggesting *S. pneumoniae* relies on Ami for threonine. However, the requirement of Ami for methionine, isoleucine, or lysine is not as significant because we did not detect genetic interactions between *ami* and genes involved in their synthesis. Methionine and isoleucine, like lysine, are probably imported via an alternative transporter(s). AmiC interacts negatively with the *spd_0751-spd_0753* operon, further confirming the latter are involved in amino acid salvage (**Fig. 3C**).

### YjbK is likely an activator of MurA

*S. pneumoniae* harbors two UDP-N-acetylglucosamine 1-carboxyvinyltransferases, MurA (Spd_1764) and MurZ (Spd_0967), that both catalyze the condensation of UDP-GlcNAc with phosphoenolpyruvate (PEP) for peptidoglycan biosynthesis ^52^. Cells lacking both genes are nonviable (**Fig. 5A to 5D**) ^52^. Unlike in most bacteria, MurZ appears to be the more active UDP-N-acetylglucosamine 1-carboxyvinyltransferase in *S. pneumoniae* ^53^. We observed a strong negative interaction between the uncharacterized gene *yjbK* (*spd_0981*) and *murZ*, but not *murA* (**Fig. 5A to 5D**), suggesting YjbK is a regulatory factor specifically for MurA.

**Fig. 5.**
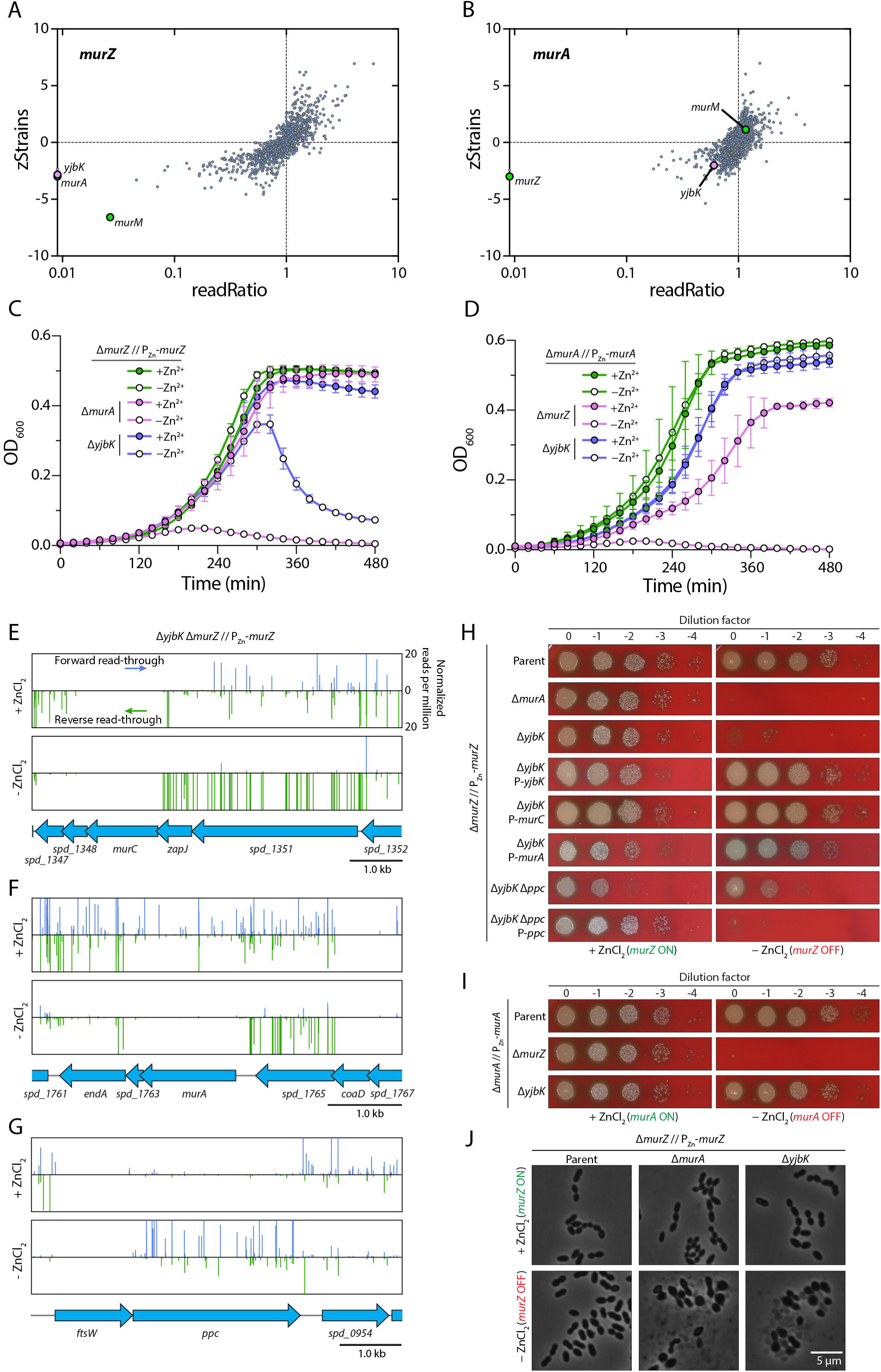
YjbK is an activator of MurA. (**A**) Dual Tn-seq revealed negative genetic interactions between *murZ* and *murA*, *yjbK* (in pink), and *murM*. (**B**) A negative genetic interaction was also detected between *murA* and *murZ* (in green). Data points on the y-axis have a readRatio value of 0. (**C**) Δ*yjbK* cells lysed when *murZ* was depleted. Strains NUS4567 [Δ*murZ*(*spd_0967*) // P_Zn_-*murZ*], NUS4585 [Δ*murZ* Δ*murA*(*spd_1764*) // P_Zn_-*murZ*], and NUS5599 [Δ*murZ* Δ*yjbK*(*spd_0981*) // P_Zn_-*murZ*] were grown in BHI medium at 37°C in 5% CO_2_ with or without the supplementation of ZnCl_2_ and MnCl_2_. Growth was monitored by measuring the optical densities of the cultures at 600nm (OD_600_) over time. Plotted are the means of three biological replicates and the standard deviation. (**D**) Δ*murZ* cells lysed when *murA* was depleted. Growth curves on strains NUS4568 [Δ*murA* // P_Zn_-*murA*], NUS4588 [Δ*murA* Δ*murZ* // P_Zn_-*murA*], and NUS5601 [Δ*murA* Δ*yjbK* // P_Zn_-*murA*] as performed above. RB Tn-seq was performed with strain NUS5599 in the presence or absence of ZnCl_2_ and MnCl_2_ supplementation. See the legends of Fig. 2 for details. Tn insertions in the *murC* (**E**), *murA* (**F**), and *ppc* (**G**) regions were shown. (**H**) Δ*murA* and Δ*yjbK* were lethal when *murZ* was depleted. The lethality was alleviated when *murA* and *murC* were expressed ectopically or when *ppc* was inactivated. The indicated strains (**Table S4**) were grown in BHI broth with added ZnCl_2_ and MnCl_2_ at 37°C in 5% CO_2_ until the OD_600_ was between 0.2 and 0.4. Cultures were normalized based on their optical density, serially diluted, and spotted on blood agar plates with (left) or without (right) added ZnCl_2_ and MnCl_2_. Plates were incubated overnight at 37°C in 5% CO_2_ before imaging. Genes following “P-” are overexpressed from a constitutive promoter. Experiments were done twice with similar results. (**I**) Δ*yjbK* was viable when *murA* was depleted. Strains NUS4568, NUS4588, and NUS5601 were grown in BHI broth with added ZnCl_2_ and MnCl_2_ at 37°C in 5% CO_2_, serially diluted, and spotted on blood agar plates with (left) or without (right) added ZnCl_2_ and MnCl_2_. Plates were incubated overnight at 37°C in 5% CO_2_ before imaging. Experiments were done twice with similar results. (**J**) Strains NUS4567, NUS4585, and NUS5599 were grown in BHI broth with or without added ZnCl_2_ and MnCl_2_ at 37°C in 5% CO_2_. Cells were imaged after the onset of cell lysis (5 hours for Δ*murA* and 6 hours for Δ*yjbK*). The parent strain (NUS4567) grown without zinc was imaged in log phase. The experiment was done twice with similar results. Bar, 5µm.

YjbK belongs to the CYTH superfamily, which is found across all domains of life ^54^. YjbK has a triphosphate tunnel metalloenzyme (TTM) fold that is thought to bind triphosphorylated substrates, but it lacks an HFXXXXEXEXK motif required for nucleotide cyclase activity ^55^. In general, the biochemical activities of CYTH superfamily proteins remain unknown ^55^. Depleting MurZ but not MurA caused the cells to deform and lyse in a Δ*yjbK* background (NUS5599; Δ*murZ* Δ*yjbK* // P_Zn_-*murZ*) (**Fig. 5I and 5J**), indicating that peptidoglycan synthesis was inhibited. We sought to identify mutations that suppress the loss of *murZ* and *yjbK* by RB Tn-seq. Consistent with a reduced MurA activity, Tn insertions that presumably led to the overexpression of *murA* or *murC* (*spd_1349*), as well as the inactivation of phosphoenolpyruvate carboxylase (Ppc, Spd_0953), alleviated the growth defect caused by Δ*yjbK* and *murZ* depletion (**Fig. 5E to 5G**). MurC installs the first L-Ala residue to UDP-MurNAc, the product of MurA/MurZ and MurB, while Ppc (Spd_0953) converts PEP, a substrate of MurA, to oxaloacetate. Thus, Δ*ppc* is expected to increase MurA activity. We confirmed that ectopic expression of *murA*, *murC*, and *yjbK*, as well as Δ*ppc*, rescued the synthetic lethality of *murZ* and *yjbK* (**Fig. 5H**). We conclude that YjbK is likely required for MurA function. This model explains why Δ*yjbK* is attenuated ^56^ since it reduces peptidoglycan synthesis. Whether YjbK modulates MurA activity by direct protein-protein interaction or indirectly by influencing substrate availability remains to be determined.

## DISCUSSION

A deeper understanding of gene function is required to find new strategies for combating the rising problem of antimicrobial resistance ^57^. The rapid advancement of next-generation sequencing technology has offered an unprecedented opportunity for functional genomics, as it enables high-throughput methods to link genotypes with phenotypes ^58^. There is a growing body of work that underpins the power of identifying gene interactions in studying gene function. For example, RodZ was discovered by isolating mutants that can grow only in conditions where the elongasome is dispensable ^59^. Other examples include the identification of the lipoteichoic acid ligase TacL ^60,61^, the capsule length control factor CpsCD ^62^, lipoproteins that control cell wall synthesis ^63,64^, and the cell division regulator KhpAB ^65^. Here, we report a novel strategy, Dual Tn-seq, that combines RB Tn-seq with the Cre-*lox* system to systematically mine genetic interactions. By evaluating the fitnesses of double mutants en masse, we identified a new CTP synthase, uncovered a regulator for cell wall synthesis, and revealed candidates involved in central metabolism.

The genetic interactions between *de novo* synthesis and salvage pathways allow us to confirm functional annotations and simultaneously identify new factors involved in metabolism. For example, the screen reported nearly all factors required for vitamin B6 synthesis (**Fig. S12**), but we have yet to find a strong candidate for the PNP/PMP oxidase. Most likely, PL in the medium alone is sufficient to support the growth of the Δ*pdxST* mutant without the need to oxidize PN and PM ^45^. This hypothesis will be tested in the future with minimal media. Similarly, we found a candidate transporter for amino acid salvaging (*spd_0751-spd_0753*) based on its negative genetic interactions with *ami* and *glnP*. Nevertheless, the Δ*glnP* Δ*glnA* double mutant appears sick (**Fig. 3B**), indicating that the alternative amino acid salvage pathways cannot sustain the wild-type growth rate. Thus, GlnPQ is the primary transporter for glutamate and glutamine in *S. pneumoniae* ^66^. Other oligopeptide import systems (*ami* and *spd_0751-spd_0753*) play relatively minor roles in growth when *glnPQ* are deleted.

The alternative CTP synthase identified in this study, PyrJ, is mostly found in Firmicutes and Actinobacteria ^31^ (**Fig. S13**). As expected, ∼31% of genomes harboring a *pyrJ* homolog lack *pyrG* (**Fig. S14**), including key pathogens like *Clostridium tetani*, *Corynebacterium diphtheriae*, and *Clostridioides difficile*. Consistently, *pyrJ* was shown to be essential in some of these species ^67,68^. Bacteria harboring PyrJ may have a competitive advantage when glutamine is limited, as PyrG activity is decreased (**Fig. 2B and 3B**). PyrJ comprises an N-terminal P-loop ATPase-like domain and a C-terminal domain with a cytidine deaminase-like fold. We speculate PyrJ emerged by coupling an ATPase with a cytidine deaminase, thus powering the reverse reaction by ATP hydrolysis. This would also explain why PyrJ is unable to use glutamine as an nitrogen donor.

Except for Clostridia, YjbK homologs are commonly found in Firmicutes (**Fig. S15**). We found that YjbK is required for MurA function. This result may explain why overexpression of MurZ, but not MurA, caused an aberrant cell shape in pneumococcus ^53^. Although they are produced at similar cellular amounts, MurZ activity seems to predominate that of MurA ^53^, suggesting that most MurA may be inactive inside the cell. Upon MurZ inactivation, YjbK may correct the defect by modulating MurA activity without changing its cellular amount ^53^. MurZ and MurA are also controlled by IreB, an allosteric inhibitor phosphorylated by StkP ^53^. IreB phosphorylation likely reduces its interaction with MurZ and MurA, thereby derepressing the system ^53^. We postulate that YjbK is a failsafe mechanism for adjusting MurA activity upon sensing a decrease in MurZ activity. Whether it directly or indirectly controls MurA awaits to be discovered. It is also intriguing why MurA in organisms outside Firmicutes do not require YjbK for function.

There are a few limitations of Dual Tn-seq. First, the distribution of insertions in the single Tn mutant libraries is intrinsically biased. This bias could be caused by Tn insertion hotspots, deletions that affect natural competence, and the fitness of the single mutants. Crossing the single mutant libraries will amplify the bias, leading to several abundant double mutants that consume considerable sequencing power. Secondly, TIS approaches are not designed for interrogating essential genes because their Tn mutants are expected to be eliminated during plating. Consequently, we expect many synthetic viable pairs to be undetectable by Dual Tn-seq. Like GIANT-coli, we also notice small blind spots (∼3 kb) in which the transformability is reduced when constructing the double mutants. Lastly, the double mutants were plated at a high density, which may result in a discrepancy in phenotype when the mutants were reconstructed.

Functional genomic approaches to systematically assay genetic interactions in bacteria hold great promise in uncovering hitherto uncharacterized genetic dark matter. Our presented Dual Tn-seq approach is one such strategy. Recently, related efforts to assay GIs in bacteria have combined CRISPR interference and TIS to probe the interactions between essential gene knockdowns and Tn mutants ^69^, or used CRISPRi strategies to knock-down the expression of multiple genes ^70,71^. The CRISPRi-based approaches have the advantage of partial knockdowns, thus permitting the analysis of essential genes, with the added burden of generating guide RNA libraries ^70,71^. Nevertheless, we envision each of these strategies to be advantageous for particular bacteria and different questions. Dual Tn-seq has the advantage that TIS methods (including RB-TnSeq) have been readily applied to many diverse bacteria, so its general implementation should be possible. Because we interrogated a bacterial genome with a smaller size (with ∼2,000 protein-coding genes), implementation in a bacterium with a larger genome will require the construction of more double mutants and increased sequencing depth. The latter should not be an issue as sequencing costs keep dropping. Given this reality, it should also be straightforward to profile the fitness of Dual Tn-seq double mutant libraries under different conditions to identify condition-specific GIs. As such, we envision Dual Tn-seq as an important tool for the high-resolution interrogation of bacterial molecular networks.

## METHODS

### Bacterial strains, plasmids, and growth conditions

The strains used in this study are listed in **Table S4**. Unless otherwise specified, *S. pneumoniae* cells were grown in brain heart infusion broth (BHI) or on tryptic soy agar plates supplemented with 5% (vol/vol) sheep blood (blood plates) (Biomed Diagnostics, 221261) at 37°C in 5% CO_2_. *Escherichia coli* was grown in lysogeny broth (LB) for most experiments or in terrific broth for protein expression. Where indicated, antibiotics were added at final concentrations of 0.3 μg/mL for erythromycin (Erm), 300 μg/mL for kanamycin (Kan), 300 μg/mL for streptomycin (Str), and 200 μg/ml for spectinomycin (Spec) for *S. pneumoniae*. For *E. coli*, ampicillin and kanamycin were added at final concentrations of 100 μg/mL and 50 μg/mL, respectively. Expression of the *czcD* promoter (P_Zn_) was induced by adding ZnCl_2_ to a final concentration of 450 μM. To counteract the cell division defects caused by Zn^2+^ toxicity, 45 μM MnCl_2_ was added to the medium simultaneously ^72,73^.

### Strain construction and transformation

Primers for generating PCR amplicons are listed in **Table S5**. PCR products were synthesized using Phusion DNA polymerase unless otherwise specified. Genetic cassettes were assembled using Gibson assembly ^74^. Pneumococcal strains were constructed by transforming cells with PCR amplicons, assembled fragments, or genomic DNA (gDNA) after inducing natural competence as described previously ^75,76^. Briefly, 1 mL of log phase culture was incubated with 1 mM CaCl_2_, 0.04% (w/v) bovine serum albumin, and 250 ng of competence stimulating peptide (CSP-1) for 10 min at 37°C in 5% CO_2_. DNA was added, and the cells were incubated for 1.5 hours at 37°C in 5% CO_2_ before being plated on blood agar supplemented by the indicated antibiotics. Markerless deletion mutants were generated using the Janus (P-*kan*-*rpsL*^+^) or Sweet Janus (P-*sacB*-*kan*-*rpsL*^+^) cassettes with counter-selections with Str and 5% (w/v) sucrose ^77,78^. gDNA was extracted using the DNeasy Blood & Tissue kit (Qiagen) as per manufacturer’s instructions.

### Measurement of growth and microscopy

For spot dilutions, strains were grown in the indicated media at 37°C in 5% CO_2_ until OD_600_ was between 0.1 and 0.5. Cultures were diluted to OD_600_ of 0.05 and ten-fold serially diluted. Two microliters of each dilution were spotted onto blood agar plates with or without added ZnCl_2_/MnCl_2_. Plates were incubated overnight before imaging. For growth curves, strains were grown in the indicated media at 37°C in 5% CO_2_ until OD_600_ was between 0.1 and 0.5. Cultures were diluted to OD_600_ of 0.01 with BHI broth to a final volume of 200 µL before being transferred to 96-well plates. If the cells were grown in broth supplemented with ZnCl_2_/MnCl_2_, they were washed twice in 1 mL of phosphate-buffered saline before diluting to OD_600_ of 0.01 with BHI broth to a final volume of 200 µL. Growth measurements were performed by incubating the plate in a Tecan Spark 10M microplate reader at 37°C. OD_600_ measurements were taken every 20 minutes after a short pulse of shaking for 20 seconds. For microscopy, cells were added to 1% (w/v) agarose pads and imaged using an Olympus IX81 inverted microscope equipped with a Hamamatsu C11440 digital camera.

### Construction of plasmid vectors comprising the RBlox-transposons

pZIK244 harboring RBloxSpec was constructed by cloning an amplicon comprising the Magellan6 Tn ^13^ generated with primers O662 and O663 using the Zero Blunt TOPO Cloning Kit (Invitrogen). These primers added the D1/D2 priming sites and the *lox71* site to the Tn. The barcoded pZIK244 was electroporated into Electromax DH10B cells (Invitrogen) following the manufacturer’s instructions.

pZIK251 harboring RBloxErm was constructed by cloning a cassette comprising a codon-optimized *ermAM* ^79^, U1 and U2 priming sites, and the *lox66* site. First, we synthesized the cassette chemically (gBlocks, IDT) and amplified it with primers O2269/O2270 to add the Mariner inverted repeats. The amplicon was cloned using the Zero Blunt TOPO Cloning Kit (Invitrogen). The intermediate vector was used as a template for PCR with primers O2377 and O2378. The resulting amplicon was DpnI-treated, assembled by Gibson assembly, and transformed into NEB 5-alpha (NEB) cells following the manufacturer’s instructions to produce pZIK251.

Barcoded transposon vector pools were generated by PCR amplification of plasmids pZIK244 and pZIK251 with primers O930 and O931 (for pZIK251) or O1078 and O1079 (for pZIK244) that add 20- nucleotide random barcodes and the overlapping regions for Gibson assembly. We used 200 ng of template plasmid per PCR to maximize barcode diversity and did 18 (for pZIK244) or 25 (for pZIK251) independent PCRs. The reactions were DpnI-treated, and the amplicons were pooled and gel extracted using the Zymoclean Gel DNA Recovery Kit (Zymo Research). Purified amplicons were ligated via Gibson assembly, purified using the DNA Clean & Concentrator kit (Zymo Research), and electroporated into Electromax DH10B cells (Invitrogen) as per manufacturer’s instructions. The RBloxErm and RBloxSpec plasmid pools have an estimated 8.8 million and 15.2 million unique barcodes, respectively.

### Library Construction

The RBloxErm and RBloxSpec plasmid pools were extracted using the QIAprep Spin Miniprep Kit (Qiagen). gDNA was extracted from NUS1927 [*rpsL1* Δ*bgaA*::*t1t2-*P_Zn_-*cre*]. Construction of Tn libraries was performed as described previously ^62^ with modifications. In vitro transposition was done by mixing 12.5 µl 4x buffer A (100 mM Tris-HCl 8.0, 400 mM NaCl, 60 mM MgCl_2_, 40% (v/v) glycerol, 8.0 mM 1,4 dithiothreitol [DTT], and 1.0 mg/mL bovine serum albumin [BSA]), 1.5 µg genomic DNA prepared from strain NUS1927, 1.5 µg barcoded RBloxSpec or RBloxErm plasmid pool, and 0.14 µg purified MarC9 transposase in 50 µL reaction volumes. The mixture was incubated for 1.5 hours at 30_J°C and purified using the DNA Clean & Concentrator kit (Zymo Research). Transposed DNA was diluted in water to a final volume of 25.5 µl, and then 3.3 µl buffer B (0.5 M Tris HCl pH 8.0, 100 mM MgCl_2_, and 10 mM DTT), 1.65 µl 20 mg/mL BSA, 1.25 µl 2 mM dNTPs, and 1.25 µl T4 DNA polymerase (NEB, M0203L) were added to the DNA and incubated for 30 min at room temperature. After incubation, 1.25 µl 2.6 mM NAD^+^ and 1.25 µl *E. coli* DNA ligase (NEB, M0205L) were added to the mixture and further incubated at room temperature overnight. The repaired DNA was transformed into strain NUS1927. Transformants were collected from the plates, pooled, and stored in 15% glycerol at -80°C. We constructed two libraries for RBloxSpec, one library (ML1) was used to generate 88 million mutants of our dual-TnSeq library, and a second library (ML3) for the remaining 1.33 billion mutants (see below).

### Transposon Library Mapping

Tagmentation of gDNA isolated from the RBlox transposon libraries was done using the Illumina DNA Prep (M) kit according to the manufacturer’s instructions with the following modifications. Tagmented DNA was amplified with custom primers O1282 and O2417 (for RBloxErm) and O1283 and O1414 (for RBloxSpec). The ’post-tagmentation cleanup’ step was done to enrich for transposon junctions and add the P7 and i7 sequences, using the following thermocycler settings: 68°C for 3 min; 98°C for 3 min; 25 cycles of 98°C for 45 sec, 62°C for 30 sec, 68°C for 2 min; and lastly 68°C for 1 min. Following the ’clean-up libraries’ steps, P5, i5, and read1 sequences were added via amplification of the product with O1282 and O1143 for RBloxErm and O1283 and O1144 for RBloxSpec using Phusion polymerase and the following settings: 95°C for 3 min; 20 cycles of 95°C for 30 sec, 60°C for 30 sec, 72°C for 1 min; and lastly 72°C for 5 min. Amplicons were sized-selected using AMPure XP Reagent (Beckman Coulter) at a 0.5x ’beads to sample ratio’ and sequenced via the Illumina platform at Azenta Life Sciences. We sequenced four preparations of the ML1 library and three preparations of the ML2 and ML3 libraries to increase the number of usable barcodes. We also mapped the library two more times using another method ^20,80^. Briefly, after shearing the DNA purified from the libraries to ∼300 bp, we used the KAPA Hyper Prep Kit to ligate a splinkerette adapter (**Table S5**), followed by an initial round of PCR to enrich for transposon insertion sites (with oligos Rd1_FOR_tnseqV3_Zik_SpecR and Rd1_REV_tnseqV3 for the RBloxSpec library, and Rd1_FOR_tnseqV3_Zik_ErmR and Rd1_REV_tnseqV3 for the RBloxErm library). After Ampure bead cleanup, we performed a second round PCR reaction with the appropriate round 2 (Rd2) oligos that contain Illumina adapter sequences and dual unique indexes (**Table S5**). After cleanup, the round 2 PCRs were sequenced on an Illumina NovaSeq instrument (2 x 150bp reads) at Novogene.

All of the above sequencing reads were combined and mapped using the Perl scripts described previously ^20^. The ML1, ML2, and ML3 libraries had 103,387; 227,041; and 620,372 unique transposon insertions with usable barcodes, respectively.

### Dual Tn-seq Library Construction

To generate the dual-TnSeq double-mutant library, frozen aliquots of the RBloxErm library in the NUS1927 background were thawed and diluted to OD_600_ of 0.05 in BHI. The diluted library was incubated for 3 hours at 37°C in 5% CO_2_. It was further diluted to OD_600_ of 0.1 in BHI and transformed with the gDNA purified from the RBloxSpec library. Typically, 200 to 500 ng of gDNA per milliliter of culture was used for generating the Dual Tn-seq libraries. Transformation reactions were incubated for 1.5 hours before being plated on blood agar with spectinomycin and erythromycin and incubated overnight at 37°C in 5% CO_2_. Transformants were collected, diluted to OD_600_ of 0.1 in BHI, and incubated for 2 hours at 37°C in 5% CO_2_. ZnCl_2_ was added to 0.85 mM to induce *cre*, and the cells were further incubated for 2 hours to allow recombination of the barcodes. Cells were collected by centrifugation at 15,000 x g for 5 min at room temperature, the supernatant was discarded, and the pellets were frozen at -80°C. We generated 1.42 billion transformants across 21 batches of independently constructed double-mutant libraries. The first 88 million of these transformants were constructed using the first RBloxSpec library (ML1) for donor gDNA, while the remaining used the ML3 library.

Cell pellets of the libraries were thawed and heat-inactivated at 85°C for 1 hour. gDNA was then extracted and quantified using the Quant-iT Qubit dsDNA HS Assay Kit (Q32851, ThermoFisher). Since each batch of the Dual Tn-seq library has variable numbers of transformants, we normalize the amount of gDNA added to the PCR based on the total number of CFUs in the library. PCR amplification was performed using Phusion polymerase with the following settings on 300 ng of combined gDNA per 50 μL reaction: 95°C for 3 min; 20 cycles of 95°C for 30 sec, 55°C for 30 sec, 72°C for 10 sec; and lastly 72°C for 5 min. The primers used incorporated P5/P7, i5/i7, and read1/read2 sequences for the Illumina platform (O1369 and O1370; O3987 and O1370). Amplicons were sequenced at Azenta Life Sciences. Collectively, we amplified and sequenced 21 double-mutant libraries across five independent NovaSeq PE150 runs.

### quantitative PCR (qPCR)

To quantify the relative amount of fused barcodes after *cre* induction, we collected ∼1 million transformants of the Dual Tn-seq library. Cells were diluted to OD_600_ of 0.1 and grown in BHI for 3.5 hours at 37°C in 5% CO_2_ before diluting the culture again to OD_600_ of 0.1 in the same medium. For each time point, cells were removed and centrifuged for 5 minutes at 15,000 x *g* at room temperature. The supernatant fraction was removed, and the pellets were stored at -80°C. Pellets were thawed and heat-inactivated by incubation at 85°C for 45 minutes. gDNA was extracted and quantified by Qubit. qPCR was performed on 1.2 ng of gDNA using the FastStart Essential Green Master kit (Roche) and primers O1635 and O1636 for the dual-barcode sequence and O1583 and O1584 for *pbp1a*. Reactions were run on a LightCycler 96 (Roche) with the following settings: 95°C for 10 min; 40 cycles of 95°C for 10 sec, 52°C for 10 sec, 72°C for 15 sec; and a final melting step of 95°C for 10 sec, 65°C for 60 sec, 97°C for 1 sec. Relative abundance was determined by the ΔΔCt method ^81^ and normalized to the copy number of *pbp1a*.

### Dual Tn-Seq Data Analysis

PEAR was used to merge Illumina paired-end reads ^82^. After extracting pairs of barcodes from each Illumina read, we filtered out pairs where each barcode had been mapped to a genomic location in the relevant RB-TnSeq library. We found that the distribution of the number of reads for each individual strain (pair of barcodes) was highly skewed, with a large minority of barcode pairs being seen just one or two times. By amplifying a limited set of defined dual-barcodes, we determined these rare barcode pairs are due to chimeric PCR. Also, we found that barcode pairs that correspond to synthetic lethal pairs of genes often had very low counts. So, strains were ignored within each run unless they had at least 3-6 reads, with varying thresholds across runs. Then, for each run, we counted the number of strains (i.e., different pairs of barcodes) and the total number of reads for each pair of genes. Only insertions within the central 10-90% of each gene were included. Since the first run had roughly 3x less strains but similar reads to the rest, we normalized this run by dividing the reads by three. Then, we summed the numbers of reads and strains across runs. Given the read and strain count for each (directed) pair, we estimated the expected numbers of reads or strains based on the proportion of reads or strains for each gene. For example, the proportion of reads for each gene pair will be:

f1 = reads for gene 1 / total

f2 = reads for gene 2 / total

f12 = f1*f2

expected12 = f12 * total reads

We then combined the actual and expected reads (or strains) in both directions to get statistics for undirected pairs. (In our data tables for undirected pairs, we always use locusId1 < locusId2.) We then adjusted these read and strain numbers for chromosomal position. Specifically, we divided the pneumococcal chromosome into 30 bins. For each pair of bins, we computed the median read ratio (actual reads / expected reads) and the median strain ratio (**Fig. S6A**). The median read ratio within each bin ranged from 0.24 to 2.9, and the median strain ratio within each bin ranged from 0.46 to 1.72. The enriched pairs of regions are near each other on the chromosome, especially near the origin or terminus of replication, or combine the origin and the terminus. These may indicate regions of the chromosome that are physically near each other, which might increase the efficiency of Cre-*lox* recombination. Given the estimated biases due to chromosomal location, we multiplied the original estimate of the expected number of reads or strains by the bias and then computed the final (adjusted) read ratio and zStrains = (strains - expected strains) / sqrt(expected strains) (**Fig. S5**). We required at least five expected double mutant strains for each gene pair to be seen for sufficient coverage. Otherwise, the gene pairs were discarded. Similarly, insertions in close proximity (within six genes) were ignored, as these are selected against upon double selection (**Fig. S6B**).

Code to analyze Illumina reads from Dual Tn-seq experiments is available from the bitbucket feba repository (https://bitbucket.org/berkeleylab/feba). barcodePairs.pl identifies pairs of barcodes in a fastq file, combineBarCodePairs.pl combines the output of barcodePairs.pl from multiple fastq files, bpFilter.pl compares the barcode pairs to the RB-TnSeq mappings and byGenePairs.pl counts strains and reads per pair of genes. R code for computing the read ratio and zStrains and adjusting for the chromosomal position is in dblStats.R (combinePairs, pairStatistics, and adjustStatistics functions).

### Estimating the false discovery rate of negative genetic interactions

To choose thresholds for zStrains and readRatio, we first shuffled the pairs of insertions that were identified by Dual-TnSeq, so that any genuine genetic interactions would be lost. Because of the variation in coverage based on position in the chromosome, for each pair, we randomly selected a new location for the second insertion among the second insertions that lie within the same bin (30th of the chromosome). The number of reads for each strain was kept the same. Given the actual data set, the shuffled data set, and thresholds for zStrains and readRatio, we can estimate a false discovery rate from the number of genetic interactions in the shuffled data set divided by the number in the actual data.

We initially chose zStrains ≤ -3 and readRatio ≤ 0.2. At these thresholds, the actual data had 320 putative negative genetic interactions and the shuffled data had 44, for an estimated false discovery rate of 44/320 = 14%. However, when we examined the putative interactions, we noticed that nine genes had eight or more interactions. These “frequent fliers” accounted for 144 of the 320 interactions (45%). We tested 12 of these gene pairs individually by constructing double mutants (**Table S2**), and none of them were synthetic lethal or yielded small colonies. Putative interactions for “frequent fliers” had milder defects (median read ratio of 0.15 instead of 0.10 for other pairs, P = 6.4 x 10^-4^, Wilcoxon rank sum test). Furthermore, while confirmed synthetic lethal and small colony-forming double mutants almost always had a functional relationship, most of the gene pairs for the “frequent fliers” did not. Of the nine “frequent fliers”, eight are clustered near the origin of replication (SPD_0004, SPD_0006, SPD_2033, SPD_2034, SPD_2051, SPD_2052, SPD_2057, and SPD_2058; SPD_2069 is the “last” gene in the genome annotation). We suspect that double mutants involving this region are obtained at lower frequencies.

After removing the interactions that involved the “frequent fliers”, we still had 176 putative interactions. We examined a random sample of 15 of these for their functional relatedness. Ten had a potential functional relationship, and five did not. Again, the pairs with functional relationships tended to have either a lower zStrains or a lower readRatio. So, besides zStrains ≤ -3 and readRatio ≤ 0.2, we decided to additionally require either zStrains ≤ -4 <u>or</u> readRatio ≤ 0.05. There are 154 genetic interactions that satisfy these thresholds if we include the “frequent fliers,” which are likely double mutants that we could not construct due to unknown reasons. As 11 “hits” are identified in the shuffled data, the false discovery rate is about 7%. If we exclude the “frequent fliers,” there will be 86 putative genetic interactions (**Table S3**). Among the pairs we tested individually, these thresholds cover 13 out of 15 synthetic lethal pairs and 3 out of 5 small colony pairs.

### RB-TnSeq analysis and BarSeq

Cultures of the indicated strains were grown until they reached the log phase (OD_600_ between 0.1 to 0.5). They were transformed with 300-500 ng of gDNA purified from the RBloxErm library and plated with appropriate antibiotics and ZnCl_2_ and MnCl_2_ where indicated. Typically, at least 1 million transformants will be obtained. Cells were collected by flooding the plates with BHI and stored at - 80°C after adding glycerol to a final concentration of 15% (v/v). For isolating suppressors, transformants were plated on blood agar plates without added ZnCl_2_ and MnCl_2_ and incubated overnight at 37°C in 5% CO_2_. Barcodes were amplified from 300 ng of gDNA purified from the surviving colonies using Phusion polymerase and the Illumina-ready primers listed in **Table S5**. For example, RBloxErm barcodes were amplified with primers O1143 and O1370 using the following thermocycler settings: 95°C for 3 min; 20 cycles of 95°C for 30 sec, 52°C for 30 sec, 72°C for 10 sec; and finally 72°C for 5 min. Amplicons were gel-purified and sequenced via the Illumina platform at Azenta Life Sciences. Insertions were quantified using Perl scripts described previously ^20^ and normalized to reads per million reads.

### Measuring transformation efficiencies

Single mutants containing an RBloxSpec or RBloxErm cassette were constructed as described above. These strains served as the template for the PCR. In general, amplicons that span 800 bp upstream and downstream of the RBloxSpec cassette were synthesized using Phusion polymerase with primers listed in **Table S5**. Amplicons were purified using the QIAquick PCR Purification Kit (Qiagen) before being transformed into recipient cells harboring RBloxErm after inducing natural competence. Briefly, 1 mL of recipient cells was diluted to an OD_600_ of 0.05, transformed with 20-45 ng of the amplicon, and placed using appropriate antibiotics. The amount of each amplicon and the recipient cultures were kept consistent for each experiment. The number of transformants was quantified as colony-forming units per milliliter. Transformation efficiency was calculated using CFU/mL using the equations outlined in **Figure S7A**.

### Construction of protein expression vectors

pZIK257 (P_T7_-*spd_0310*-*his6*) was constructed by Gibson assembly of the amplicons synthesized using primers O5132 and O5133 (for the pET-21^+^ backbone (Novagen)), and O5134 and O5127 (for *spd_0310* in D39W). The amplicon containing the pET-21^+^ backbone was pretreated with DpnI (NEB).

The assembled plasmid was introduced into DH5a(lpir), purified using the QIAprep Spin Miniprep Kit (Qiagen), and confirmed by Sanger sequencing. pZIK259 (P_T7_-*pyrG*(*spd_0442*)-*his6*) was constructed by Gibson assembly of the amplicons synthesized using primers O5220 and O5133 (for the pET-21^+^ backbone), and O5221 and O5219 (for *pyrG*(*spd_0310*) in D39W). The amplicon containing the pET-21^+^ backbone was pretreated with DpnI (NEB). The assembled plasmid was introduced into DH5a(lpir), purified using the QIAprep Spin Miniprep Kit (Qiagen), and confirmed by Sanger sequencing.

pZIK257 and pZIK259 were introduced into BL21(λDE3) using the methods described in Chung et al.^83^.

### Protein Purification

SPD_0310-His6 and PyrG-His6 were purified from strains BL21(λDE3)/pZIK257 and BL21(λDE3)/pZIK259. Briefly, cells were grown in terrific broth with ampicillin at 37°C with shaking until the culture reached the mid-log phase. Cells were induced by adding IPTG to a final concentration of 0.5 mM and harvested after 3 hours of incubation at 37°C with shaking by centrifugation at 10,000 x *g* for 10 minutes at 4°C. Pellets were stored at -80°C. They were thawed and resuspended in lysis buffer (20 mM Tris-HCl pH 7.0, 150 mM NaCl, 10 mM MgCl_2,_ and 1:1000 of the protease inhibitor cocktail (539134; Calbiochem)). Cells were lysed with two passages through a French pressure cell at 18,000 psi. Following lysis, samples were maintained at 4°C. Lysates were clarified via centrifugation at 16,000 x *g* for 30 minutes at 4°C before being further cleared via filtration through a 0.22µm filter. The supernatant was incubated with 0.5 mL of TALON resin (Takara Bio) in lysis buffer with 20 mM imidazole for 1 hour. Bound protein was washed three times via gravity filtration with lysis buffer supplemented with 40 mM imidazole, before being eluted in the same buffer with 300 mM imidazole. Purified proteins were exchanged into storage buffer (20 mM Tris-HCl pH 7.5, 50 mM NaCl, 10 mM MgCl_2_, 10% glycerol) using PD-10 desalting columns (Cytiva) before being flash-frozen in liquid nitrogen and stored at -80°C. Purification was assessed via SDS-PAGE, and protein amount was quantified via Bradford assay.

#### CTP Synthase Assay

The CTP synthase assay was performed essentially as described in ^29^. Briefly, 0.5 mM of UTP, UDP, UMP, or uridine was mixed with 0.1 mM GTP and 0.5 mM ATP in a total volume of 100µl of the reaction buffer (20 mM Tris-HCl pH 8.0, 10 mM MgCl_2_). When indicated, L-glutamine or NH_4_Cl was added to a final concentration of 2 mM and 10 mM, respectively. Next, SPD_0310-His6 or PyrG-His6 was added to a final concentration of 1 µM, and the mixture in a 96-well plate was transferred immediately to a Tecan Spark 10M microplate reader for A291 nm measurements every 1 minute at 25°C. CTP concentration was determined using the Beer-Lambert law with the extinction coefficient ε of 1520 M^-1^ cm^-1^ ^29^.

## ACKNOWLEDGEMENTS

We thank members of the L.-T.S. laboratory for helpful discussions. This work was supported by grants from the Singapore National Research Foundation (NRFF11-2019-0005), the National Research Medical Council (OFIRG23jan-0057), and the Ministry of Education (MOE-T2EP30220- 0012) to L.-T.S.. Work by M.N.P., A.P.A., and A.M.D. was supported by ENIGMA-Ecosystems and Networks Integrated with Genes and Molecular Assemblies (http://enigma.lbl.gov), a Science Focus Area Program at Lawrence Berkeley National Laboratory, and is based upon work supported by the U.S. Department of Energy, Office of Science, Office of Biological and Environmental Research, under contract number DE-AC02-05CH11231.

## AUTHOR CONTRIBUTIONS

J.J.Z. and L.-T.S. conceived the study and designed the Dual Tn-seq; J.J.Z., M.N.P., and A.M.D. conducted the experiments; A.P.A., A.M.D., and L.-T.S. acquired funding to support the work; J.J.Z. and M.N.P. analyzed data with contributions from A.P.A., A.M.D., and L.-T.S.; J.J.Z. and L.-T.S. wrote the manuscript with contributions from M.N.P., A.P.A., and A.M.D.; A.P.A., A.M.D., and L.-T.S. supervised the work; All authors read and approved the final manuscript.

