## Supplemental figure legends for "Dual transposon sequencing (Dual Tn-seq) to probe genome-wide genetic interactions"

**Figure S1. RBlox-Tn design and function.** We constructed two transposons (RBloxSpec and RBloxErm) with either a spectinomycin or erythromycin resistance cassette flanked on opposite ends by universal priming sites (D1/D2 or U1/U2), *loxP*-derivative sequences, and 20-nucleotide random barcodes (SpecBC or ErmBC). We placed the Cre recombinase under the control of a zinc-inducible promoter (P_Zn_) at a neutral locus. Upon *cre* induction, strains harboring both RBloxSpec and RBloxErm with *lox* sites in a reverse (A) or the same (B) orientation result in an inversion or splitting of the chromosome, respectively. In both cases, the final dual-barcode structure will be preserved as a short 111 bp sequence (U1-ErmBC-*lox72*-SpecBC-D2), that can be amplified as an Illumina-ready amplicon (e.g. including, but not limited to, P5/P7 and read1 sequences).

**Figure S2. Dual-TnSeq achieves cross-genome recombination without impairing strain fitness before induction.** (A) Upon *cre* induction, recombination between distantly located RBlox-transposons can be detected by PCR amplification across the recombined junction using primer sets specific to the same (P1/P2) or reverse (P1/P3) orientations. (B) The indicated strains were grown in BHI broth at 37^o^C in 5% CO_2_, induced with ZnCl_2_ at OD_600_ = 0.1, and incubated for one hour before PCR was performed on whole cells using the indicated primer sets. (C) Sanger sequencing of the recombined barcodes from amplicons in (B) revealed the intended dual-barcode sequence. (D) The presence of P_Zn_-*cre* does not impact strain fitness when not induced. The indicated strains (**Table S4**) were grown in BHI broth at 37^o^C in 5% CO_2_ until the OD_600_ was between 0.2 to 0.4 before being normalized by the optical density and released into fresh BHI broth without ZnCl_2_. Growth was monitored by measuring the optical densities of the cultures at 600nm (OD_600_) over time. Plotted are the means from three biological replicates and the standard deviation.

**Figure S3. Bar-seq on transformed RBlox-Tn libraries recapitulates known synthetic lethal relationships.** (A) RB Tn-seq assigns unique barcodes to individual insertion sites after an initial mapping step. Later, amplification of only the barcode sequences (Bar-seq) will reveal the identity of Tn insertions in a population. (B) gDNA extracted from the mapped RBloxErm library was transformed into NUS1927 [P_Zn_-*cre*] and into NUS3358 [∆*pbp1a* // P_Zn_-*cre*] (shown), NUS3359 [∆*pbp2a* // P_Zn_-*cre*], or NUS3360 [∆*macP* // P_Zn_-*cre*]. Recovered transformants were identified and quantified via Bar-seq. (C)The normalized number of Tn insertions identified from (B) over the *pbp1a*, *pbp2a*, and *macP* loci. Compared to wild-type, insertions in *pbp1a* are reduced in the Δ*pbp2a* and Δ*macP* strains, while insertions in *pbp2a* and *macP* are reduced in the Δ*pbp1a* strain.

**Figure S4. Optimizing Dual Tn-seq in *S. pneumoniae*.** (A) Varying amounts of purified RBloxSpec gDNA were used to transform 1 mL of the RBloxErm library at OD_600_ = 0.1, and the number of total transformants per mL was quantified. Each data point represents the mean ± SD of at least two replicates. (B) Violin plot of data from (A) plotted as the number of colonies recovered from 1 mL of transformation reaction, normalized to ng of transformed gDNA. (C) A lawn of transformants recovered from reactions as performed in (A) were pooled in BHI broth, diluted to OD_600_ = 0.1 in BHI, and allowed to recover for 3.5 hours at 37^o^C in 5% CO_2_ before diluting again to OD_600_ = 0.1 and inducing *cre* with 0.85 mM ZnCl_2_ and grown at 37^o^C in 5% CO_2_. qPCR was performed on gDNA extracted from cells collected immediately before and after induction at the indicated time points, using primers that amplify across the recombined dual-barcode sequence. Relative quantity (mean ± SD) from duplicate trials was determined via the ΔΔCt method normalized to the copy number of *pbp1a*.

**Figure S5. A detailed description of readRatio and zStrains calculations.**

The probability of insertions being from any single gene, based on the number of barcode reads (top-left) or strains (top-right) in the population, are used to calculate the respective expected values of dual-reads or double-mutant strains for all gene pairs. We use the variable n to refer to either the sum normalized reads or the sum number of strains, respectively. These are first calculated for the pair in one orientation (ErmBC,gene1 > SpecBC,gene2) and combined with complimentary orientation (ErmBC,gene2 > SpecBC,gene1) in step 3. The values are then adjusted using a ratio that incorporates Cre-recombination efficiency across chromosomal position and distance (see methods and **Fig S6**). All gene pairs with insufficient strain coverage (< 5) at the end of step 4 are discarded (even if expected reads are being calculated). Lastly, the actual (observed) number of dual-reads or double-mutants is compared to the expected value, and gene pairs that deviate significantly are identified.

**Figure S6. Chromosomal position and distance between RBlox-Tn’s affects Cre-recombination at a predictable rate.** (A) The chromosome was split into 30 bins, from which each pair of genes is placed in a subsequent bin corresponding to total number of bin combinations (465). For each of the 465 bins, the median ratio of observed values divided by expected values for both dual-barcode reads (left) and double mutant strains (right) are computed. These ratios are used to adjust the expected values as outlined in **Table S1** in the final analysis. (B) The distance between two insertions was binned into 1 kb intervals, and the median ratio of observed values divided by the expected values for double mutant strains (left) and dual-barcode reads (right) was calculated for each bin. We observed a higher median ratio for insertions that were near each other on the chromosome, as well as for origin x terminus pairs. We speculate that origin x terminus pairs recombine efficiently because the origin and terminus are near each other in 3D space, even though they are not nearby in the sequence.

**Figure S7. A transformation efficiency assay identifies synthetic lethal interactions.** (A) A transformation efficiency assay was used to assess double-mutant strain viability using gene pairs of varying readRatio and zStrains values. The test transformation efficiency (top ratio) measures the number of transformants per mL following the transformation of one knockout construct into the corresponding pair deletion background. To correct for strain transformability, this value is divided by the value obtained for the transformation of a neutral *ply* knockout construct. The control transformation efficiency (bottom ratio) controls the viability of the transformed cassette into a neutral background. (B) The test and control transformation efficiencies for 109 gene pairs were plotted against each other (**Table S2**). We defined synthetic lethal interactions as having >100-fold decrease in test over control transformation efficiencies. Data points flush on the x-axis gave a test value of 0. We noticed 5 resultant double-mutants that gave a small colony phenotype (color-coded in blue) compared to either parent background. (C) Spot dilutions confirm the small colony phenotypes. The indicated strains (**Table S4**) were grown in BHI broth at 37^o^C in 5% CO_2_ until the OD_600_ was between 0.2 and 0.4. Cultures were normalized based on their optical density, serially diluted, and spotted on blood agar plates. Plates were incubated overnight at 37^o^C in 5% CO_2_ before imaging. (D) Corresponding zStrains (left) and readRatios (right) from the dual-TnSeq dataset (**Table S1**) for the gene pairs assayed here and grouped based on phenotype (**Table S2**). Error bars represent mean ± SD for each category. One-way ANOVA followed by Dunnet’s test with comparison to the viable category was used to determine significance. ****p < 0.0001; ***0.0001 < p < 0.001; *0.01 < p < 0.05; n.s., not significant. (E) Corresponding zStrains and readRatios from dual-TnSeq for the gene pairs assayed here were plotted against each other. Data points flush on the y-axis have a readRatio value of 0.

**Figure S8. Converging pathways for de novo biosynthesis and salvage of pyrimidine nucleotides in *S. pneumoniae*.** Pathways leading to cytidine triphosphate (CTP) production in *S. pneumoniae*, based on annotations and on the results from this study. De novo synthesis of CTP invariably proceeds from the amination of uridine triphosphate (UTP). Alternatively, the precursor nucleosides can be salvaged and phosphorylated once in the cytoplasm. The nucleoside transporter PnrABCD prefers purine nucleosides but can import sufficient cytidine (CR) when overexpressed. Spd_1425 is an uncharacterized major facilitator superfamily (MFS) protein that we show imports at least CR. Spd_0310 (renamed PyrJ) belongs to a formerly uncharacterized protein family (DUF1846) and is shown here to be a novel ammonia-dependent CTP synthase. Other acronyms: UR, uridine; UMP, uridine monophosphate; UDP, uridine diphosphate; CMP, cytidine monophosphate; CDP, cytidine diphosphate; OMP, orotidine monophosphate.

**Figure S9. Exogenous cytidine rescue of** ∆***spd_0310* and** ∆***pyrG* synthetic lethality is abolished in nucleoside transporter mutants.** (A) NUS4797 was grown in BHI broth at 37^o^C in 5% CO_2_ with added ZnCl_2_/MnCl_2_, washed, and released in BHI broth with or without ZnCl_2_/MnCl_2_ at 37^o^C. Where indicated, cultures were further supplemented with 50 µM of either adenosine (AR), guanosine (GR), or uridine (UR). Growth was monitored by measuring the optical densities of the cultures at 600nm (OD_600_) over time. Plotted are the means from three biological replicates and the standard deviation. (B) Growth curves as performed in (A) but supplemented with all of AR, GR, and UR with or without added cytidine (CR). (C) Growth curves as performed in (A) using NUS4797 and NUS5294. Reduced growth in strains lacking *spd_1425* and *pnrABCD* highlights the contribution of nucleoside salvage towards growth. (D) Growth curves as performed in (A) using NUS5294 but supplemented with 50 µM or 500 µM of CR when indicated to demonstrate the absence of CR uptake in this background. (E) The indicated strains (**Table S4**) were grown in BHI broth at 37^o^C in 5% CO_2_ until the OD_600_ was between 0.2 to 0.4 before being normalized by the optical density and released into fresh BHI broth with 150 µM (left) or 0 µM (right) of the toxic cytidine analog 5-fluorocytidine (5FC). Growth was monitored by measuring the optical densities of the cultures at 600nm (OD_600_) over time. Genes following “P-” are overexpressed from a constitutive promoter. Plotted are the means from three biological replicates and the standard deviation. (F) Growth curves as performed in (E) on the indicated strains (**Table S4**) supplemented with 250 µM (left) or 0 µM (right) of 5FC. The growth of strains lacking *pyrG* and unable to salvage CR suggests the presence of another de novo source of CTP.

**Figure S10. Spd_0310 and PyrG are CTP synthases.** (A) 5 µL of purified Spd_0310-his6 and PyrG-his6 aliquots were separated on a 10% SDS-PAGE gel and stained with Coomassie. (B) Spectrophotometric activity assays for detecting PyrG- or Spd_0310-dependent amination of the uracil ring structure via an increase in absorbance at 291 nm. Reactions included purified PyrG or Spd_0310; 0.5 mM of either uridine triphosphate (UTP), uridine diphosphate (UDP), uridine monophosphate (UMP) or uridine (UR) as substrates; and either L-glutamine (L-Gln) or ammonia as nitrogen donors. Error bars represent the mean ± SD of three independent replicates.

**Figure S11. Redundant transporters revealed by Dual Tn-seq.** (A) *trkH* exhibited strong negative genetic interactions with both components of the redundant potassium transporter encoded by *spd_0429* and *spd_0430*. (B) The reverse analysis using *spd_0429* showed similar interactions with both *trkH* and its positive regulator *cabP*. (C) A representative gene of the inorganic orthophosphate transporter operon (*pstC2*) showed negative interactions with all five genes of the redundant *pst1* operon. (D) The reverse analysis using *pstC1* revealed interactions with all six genes of the *pst2* operon. Data points flush on the y-axis for all panels have a readRatio value of 0.

**Figure S12. Genetic interactions identified between pyridoxal phosphate salvage and *de novo* biosynthesis pathways.** (A) The pyridoxal phosphate biosynthetic and salvage pathways in *S. pneumoniae*, based both on annotations and on functional data via dual-TnSeq. G3P, D-glyceraldehyde 3-phosphate; DHAP, dihydroxyacetone phosphate; R5P, D-ribose-5-phosphate; Ru5P, D-ribulose-5-phosphate; L-Gln, L-glutamine; PL, pyridoxal; PM, pyridoxamine; PN, pyridoxine; PLP, pyridoxal 5’-phosphate; PMP, pyridoxamine 5’-phosphate; PNP, pyridoxine 5’-phosphate. (B) The PL/PM/PN transporter PdxU2 shows strong negative interactions with the PLP de novo synthesis genes *pdxR*, *pdxS*, and *pdxT*. (C) Similarly, negative interactions with all de novo synthesis genes are found with the other known salvage gene, *pdxK*, encoding pyridoxal kinase. Data points flush on the y-axis have a readRatio value of 0. (D) The PLP biosynthetic gene *pdxT* shows strong negative interactions with the salvage genes *pdxU2* and *pdxK*. (E) Similarly, negative interactions with the salvage genes are found with the transcriptional activator for de novo biosynthesis, *pdxR*.

**Figure S13. *pyrJ* is found predominantly in Gram-positive bacteria.** Phylogenetic distribution of *pyrG* (red ticks) and *pyrJ* (blue ticks) across 19,338 bacterial and archaeal reference genomes (RefSeq). We used BLASTp to search the above genomes for homologs of D39W PyrG (Spd_0442) and PyrJ (Spd_0310) using *e*-value cutoff of 10^-4^. The sequences of PyrG and PyrJ from D39W were PhyLoT (phylot.biobyte.de) was used to construct the tree and iToL (itol.embl.de) was used for visualization. *pyrG* is ubiquitous amongst bacterial and archaeal genomes, while *pyrJ* is found primarily in clusters amongst Firmicutes and Actinobacteria.

**Figure S14. Diverse bacteria encode *pyrJ* as the sole CTP synthase.** A pruned phylogenetic tree highlighting the presence or absence of *pyrG* (red boxes) in 55 bacteria that encode *pyrJ* (blue boxes). Species names are indicated followed by taxon ID (NCBI). Among genomes encoding *pyrJ*, the fraction that do not encode *pyrG* roughly follows the percentage indicated in the text (~31%). PhyLoT (phylot.biobyte.de) was used to construct the tree and iToL (itol.embl.de) was used for visualization.

**Figure S15. *yjbK* is found exclusively in Firmicutes.** Phylogenetic distribution of *yjbK* (blue ticks) across 19,338 bacterial and archaeal reference genomes (RefSeq). We used BLASTp to search the above genomes for homologs of D39W YjbK (Spd_0981) using *e*-value cutoff of 10^-4^. PhyLoT (phylot.biobyte.de) was used to construct the tree and iToL (itol.embl.de) was used for visualization. *yjbK* is widespread in most Firmicutes except Clostridia.
