## Supplementary figures and images for "Dual transposon sequencing (Dual Tn-seq) to probe genome-wide genetic interactions"

### Fig. S1

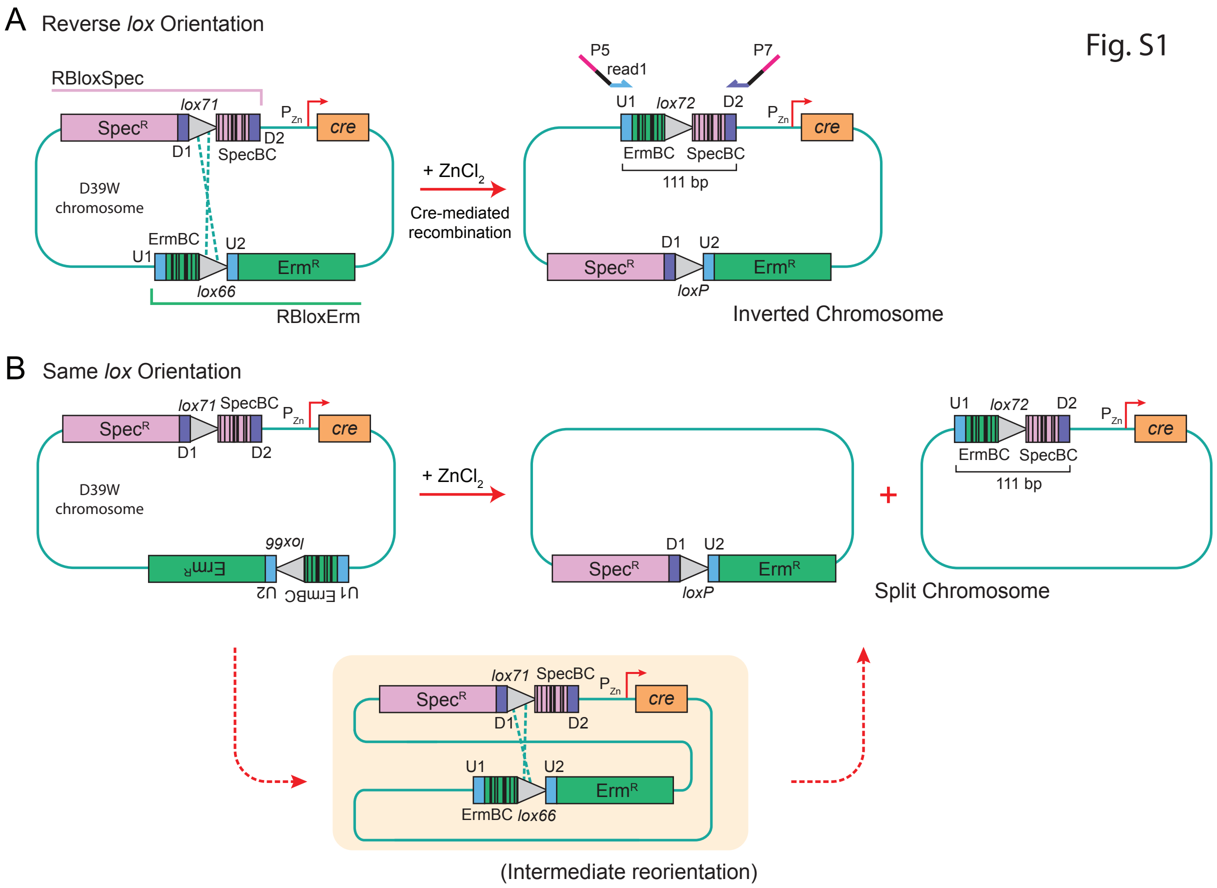

### Fig. S2

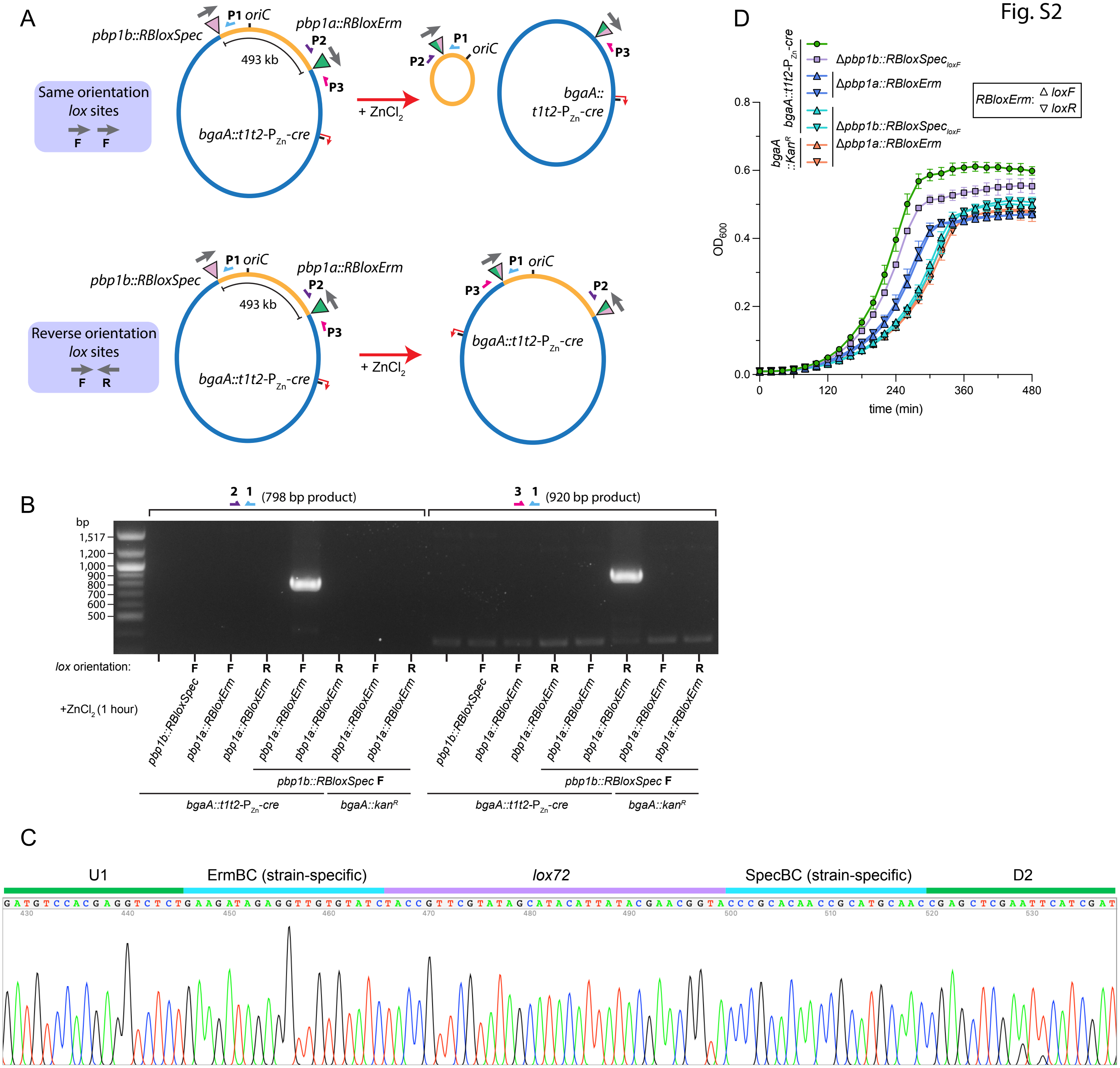

### Fig. S3

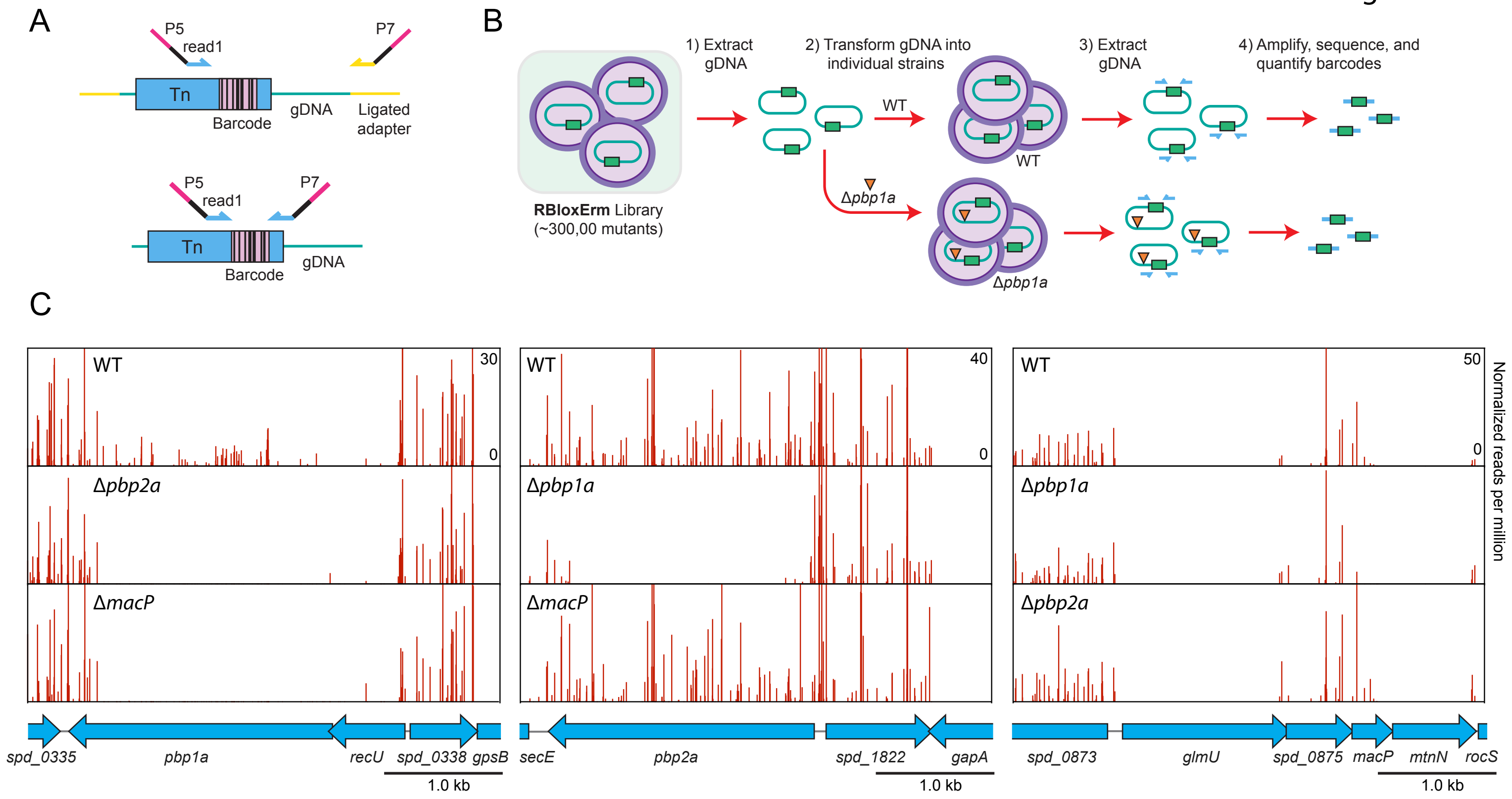

### Fig. S4

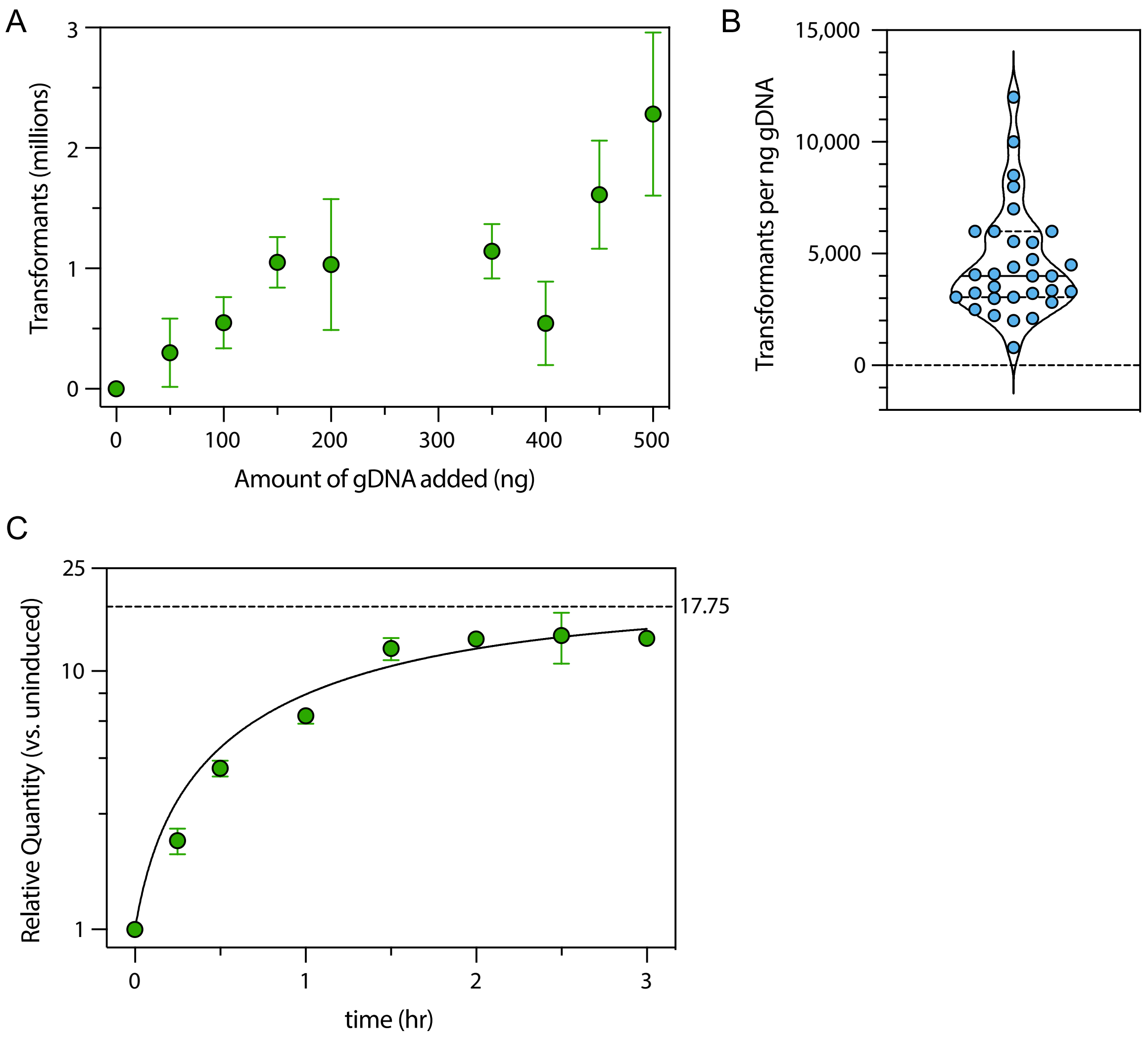

### Fig. S5

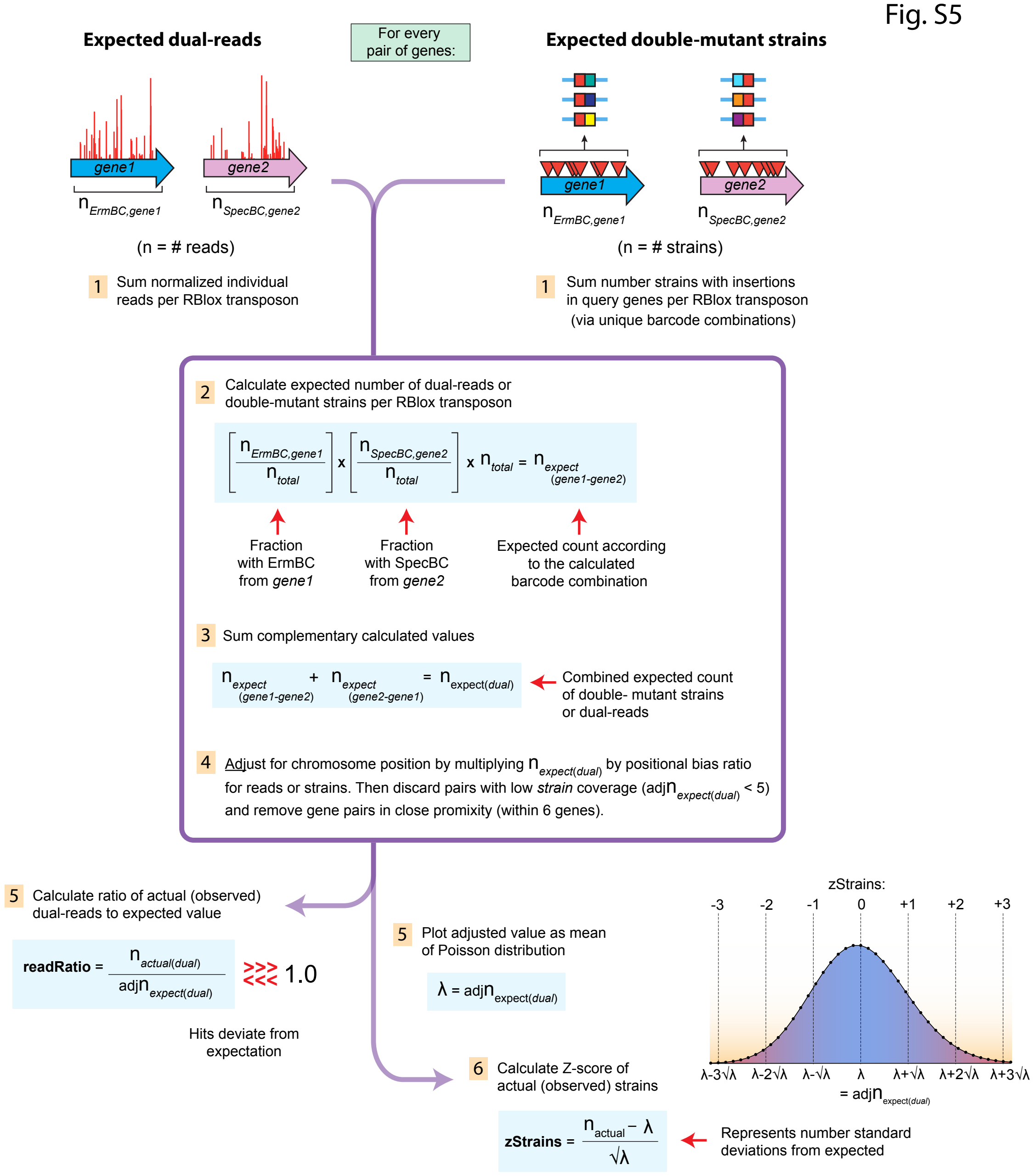

### Fig. S6

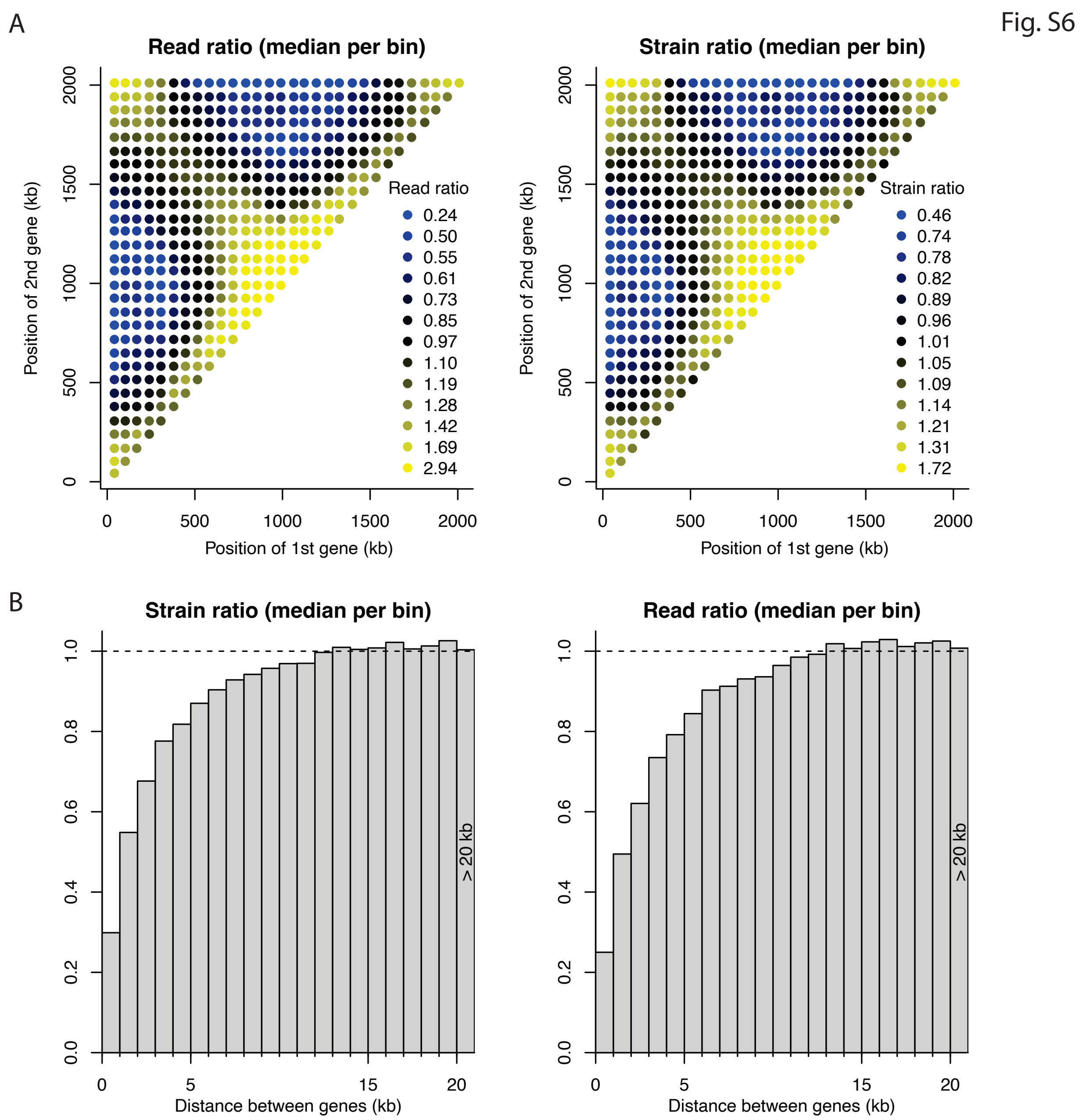

### Fig. S7

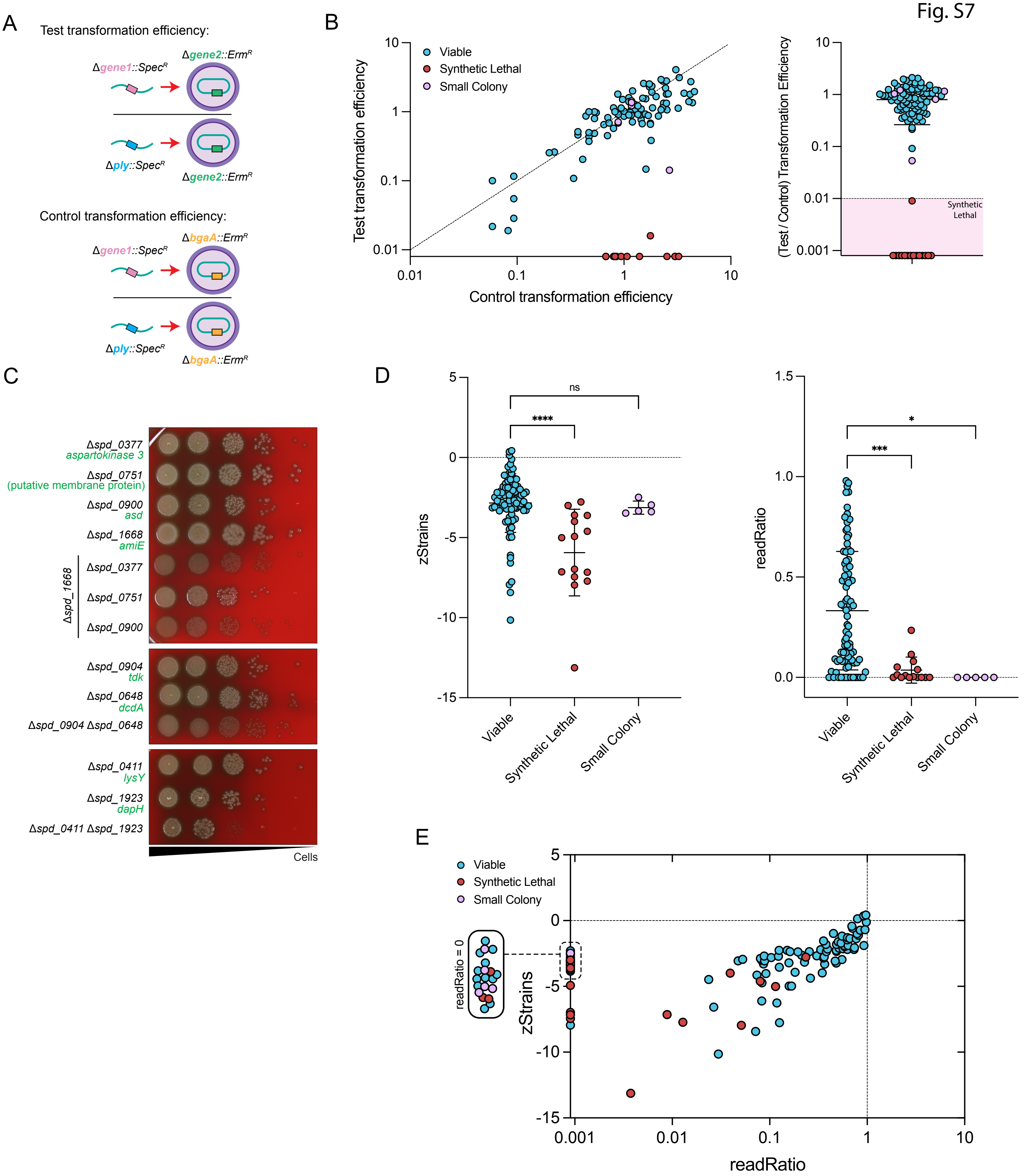

### Fig. S8

#### Fig. S8

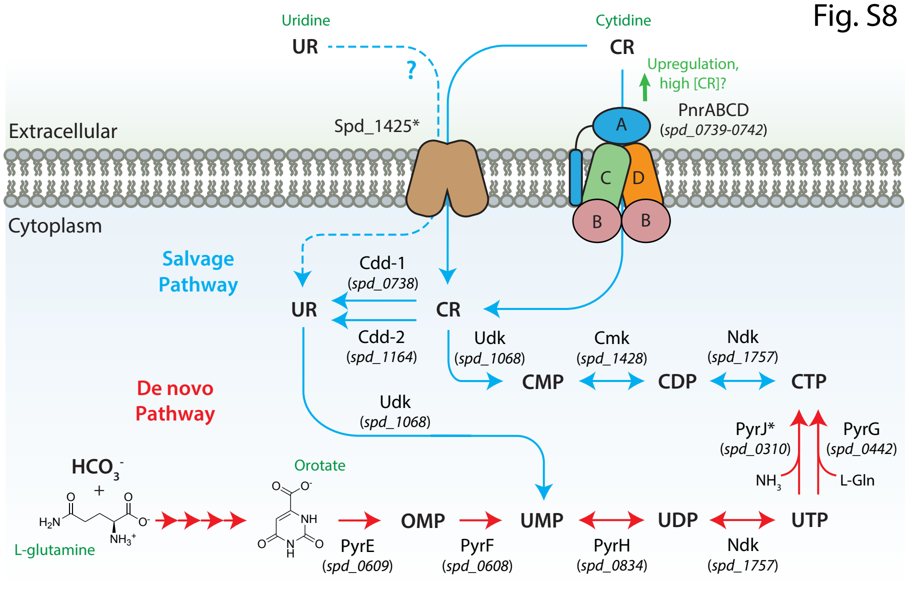


*: identified in this study

### Fig. S9

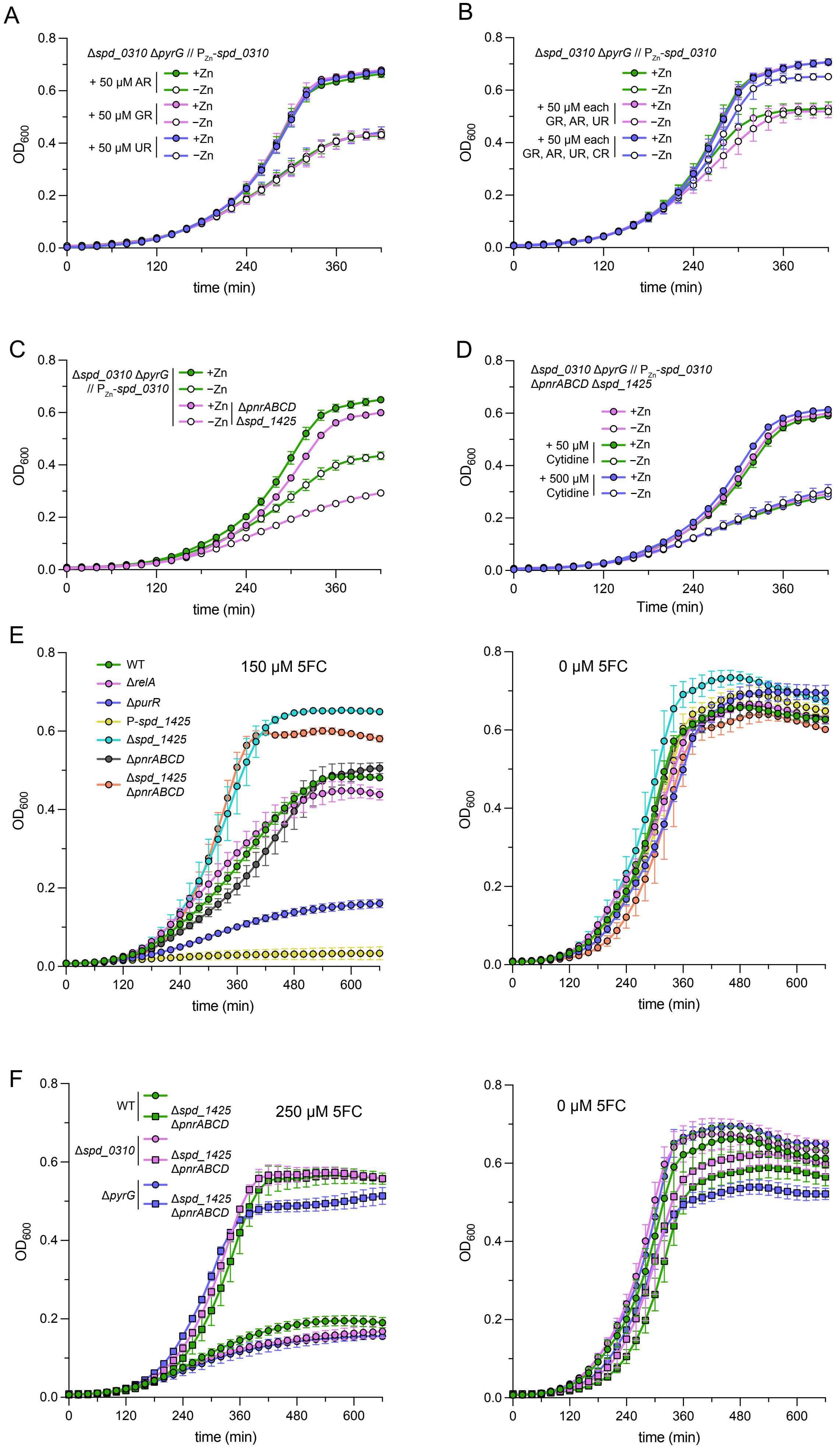

### Fig. S10

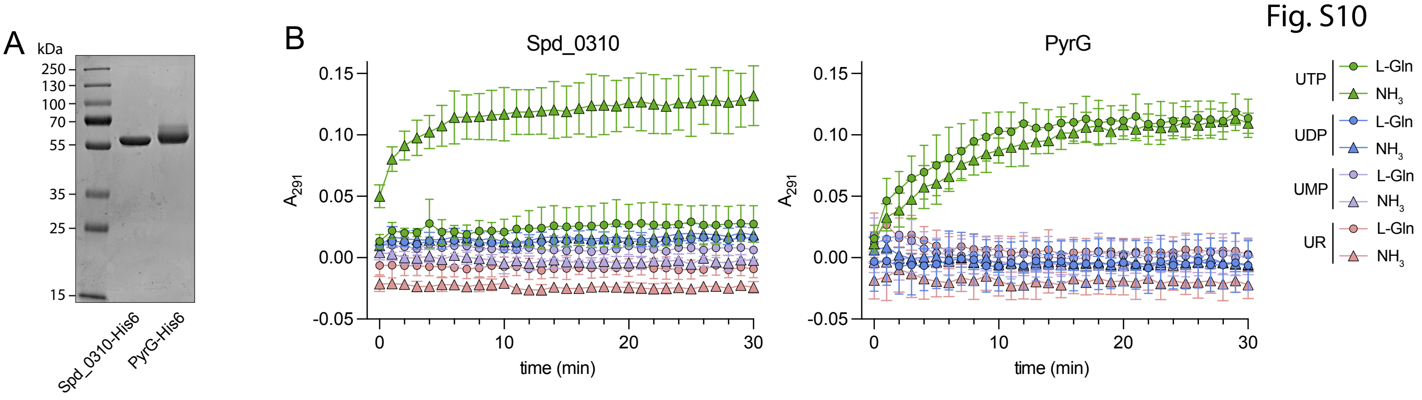

### Fig. S11

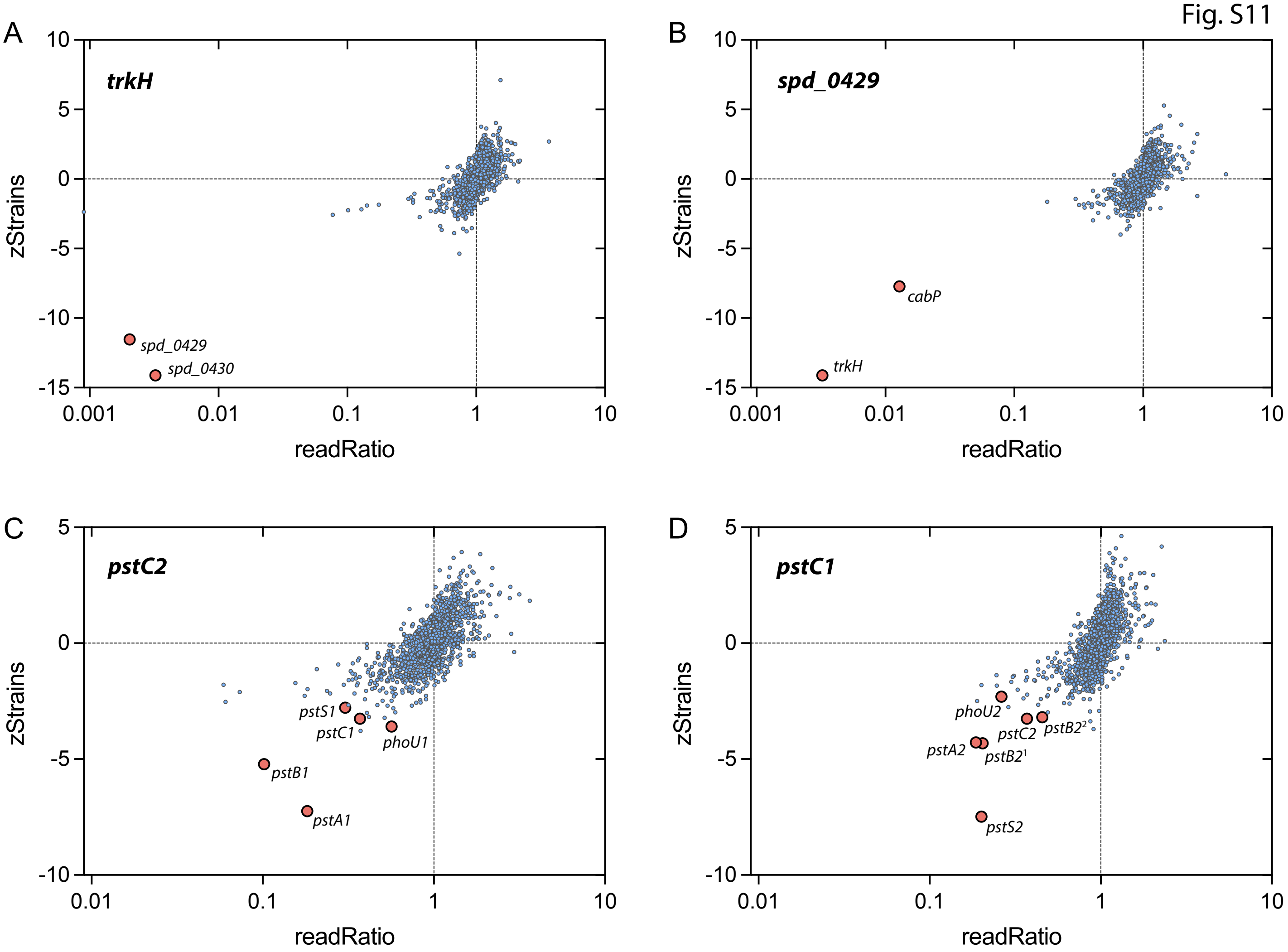

### Fig. S12

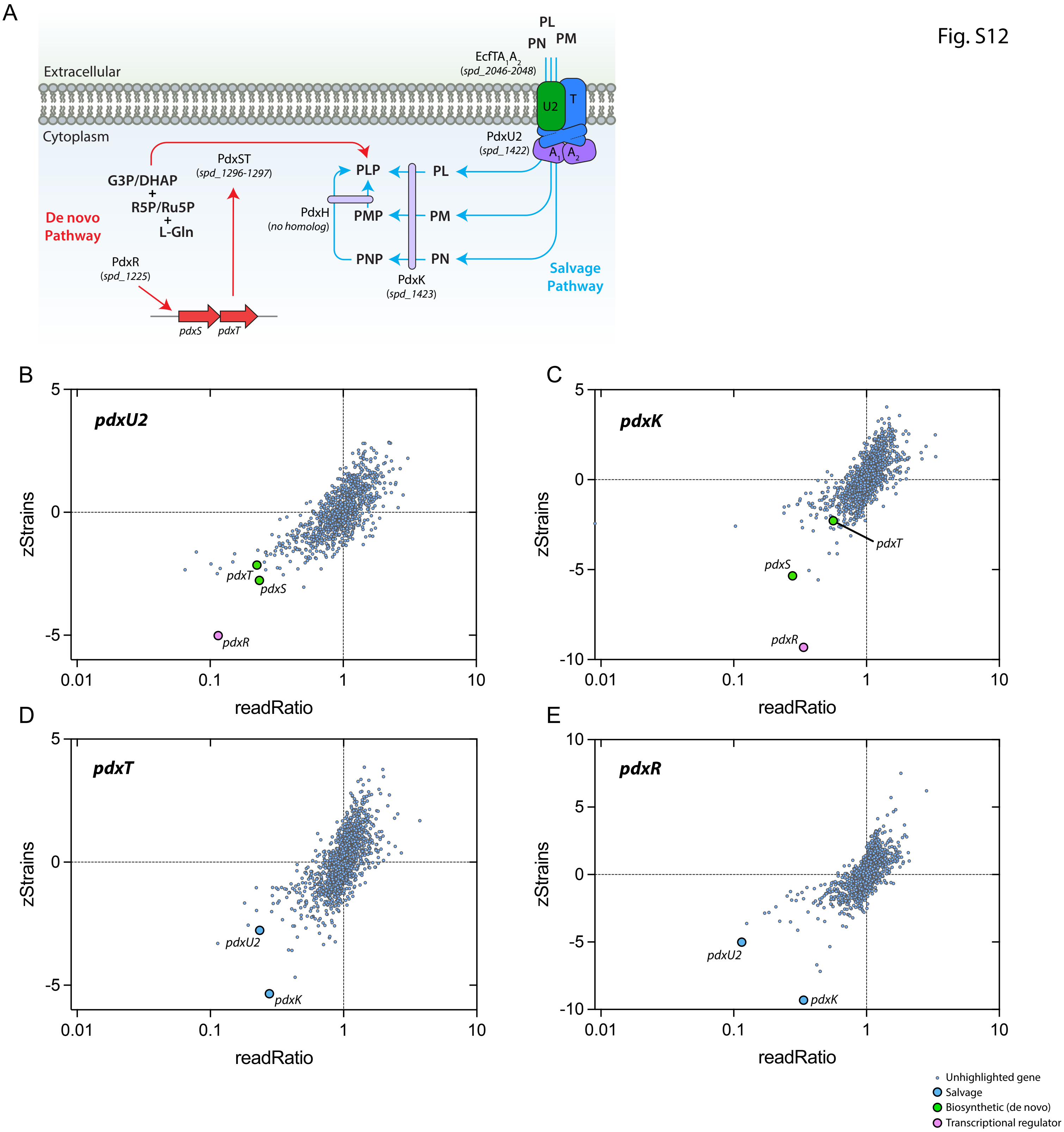

### Fig. S13

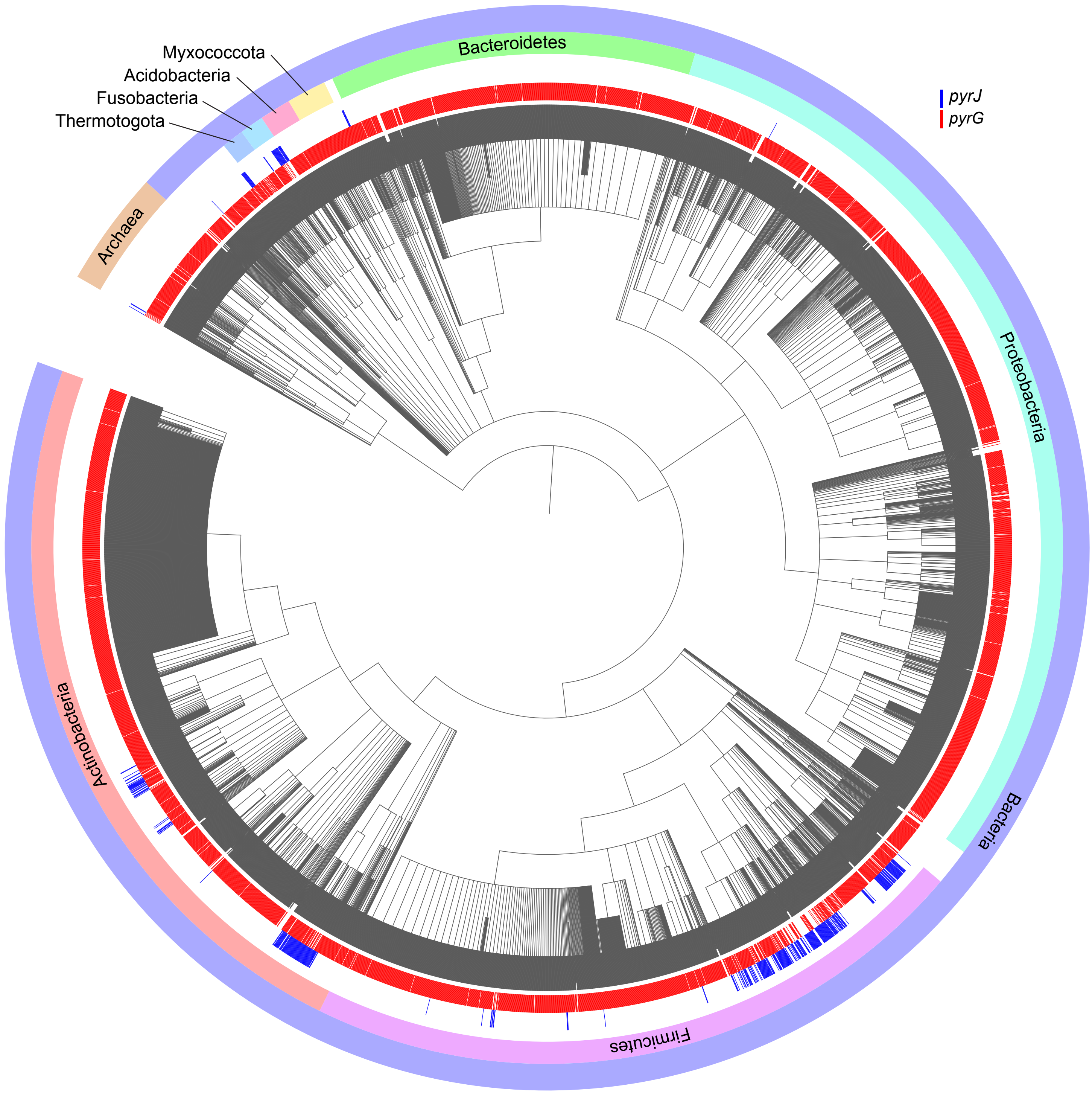

### Fig. S14

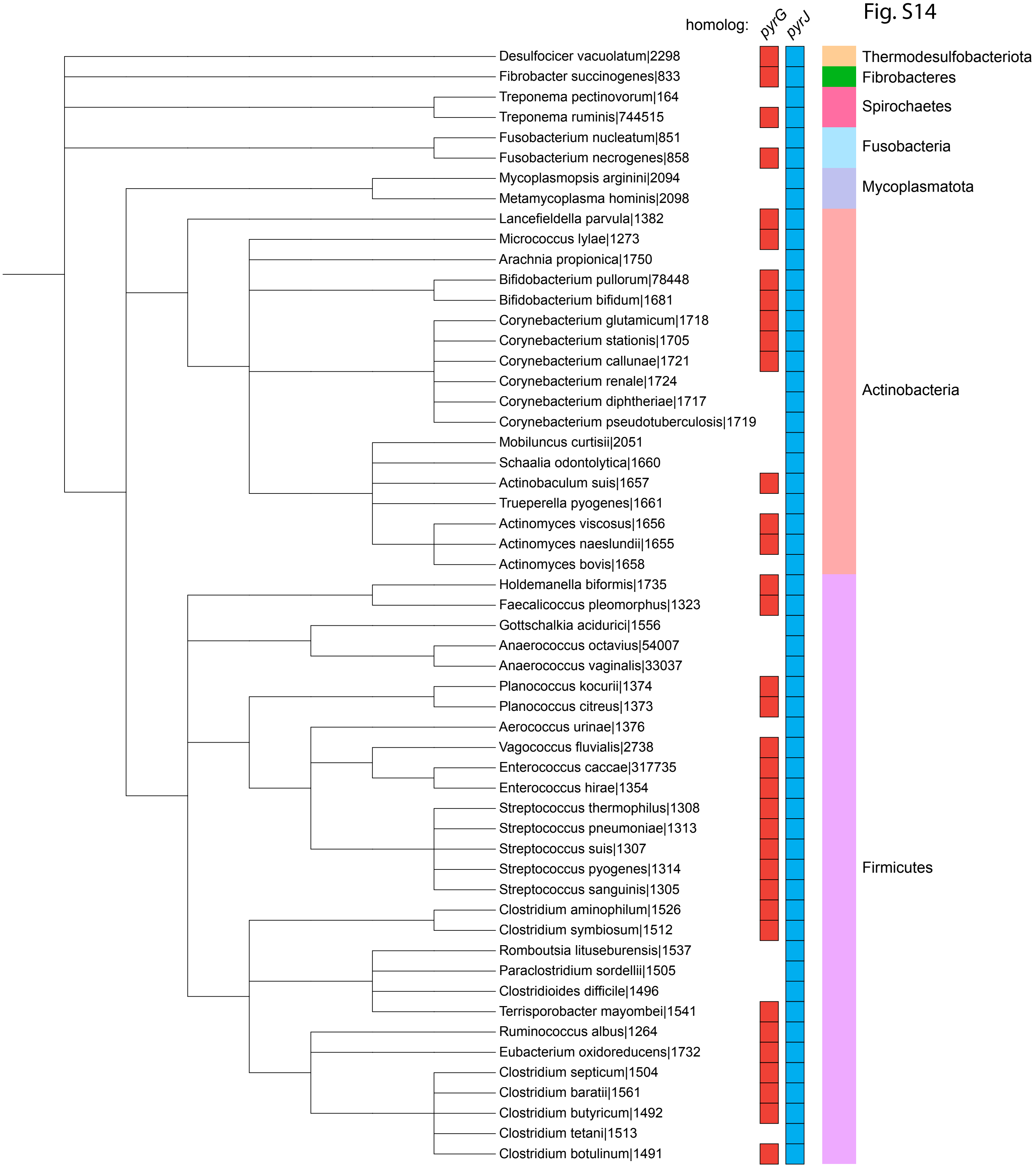

### Fig. S15

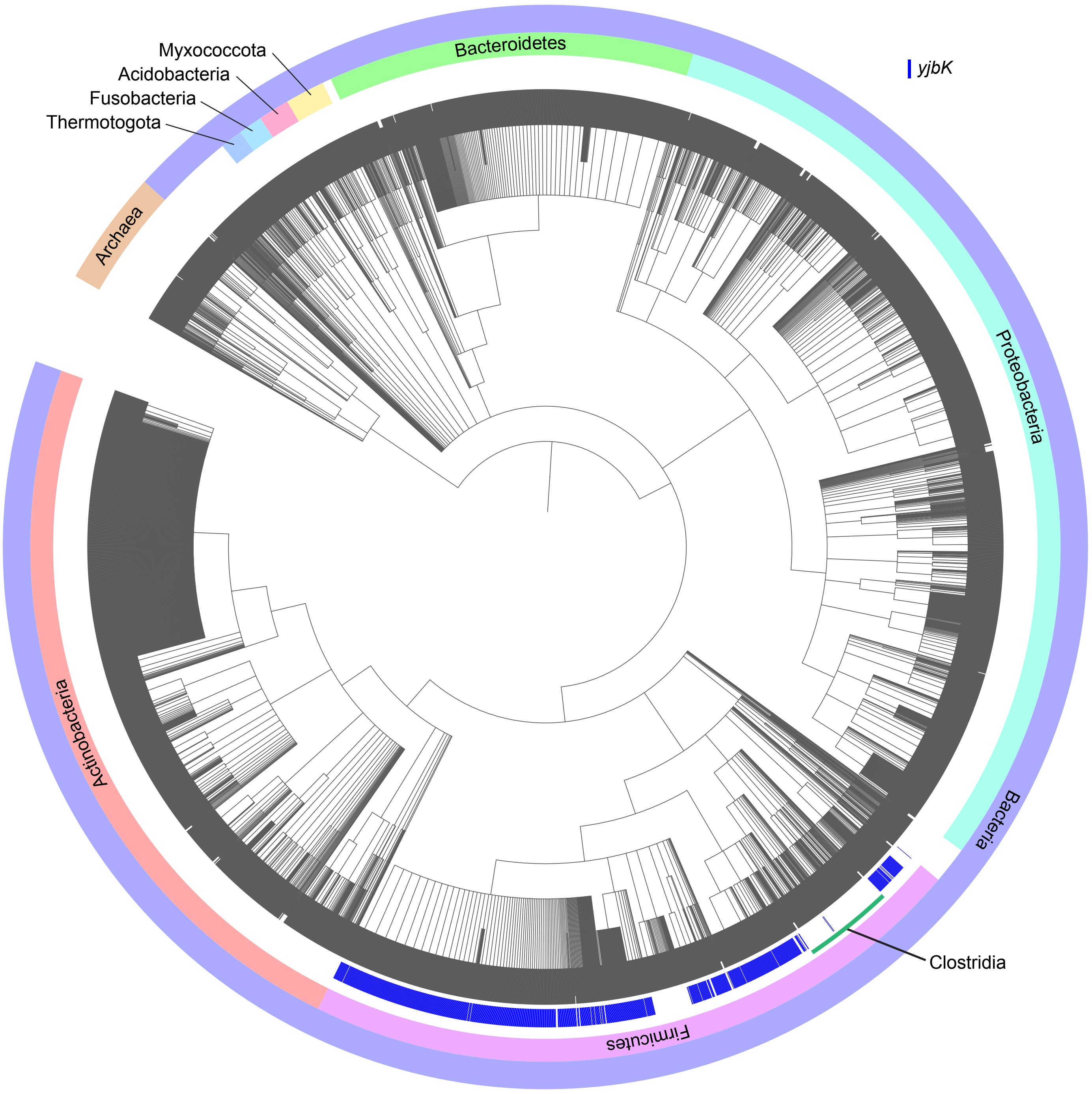
